# Thalamo-accumbal circuit adaptations following extended oxycodone abstinence

**DOI:** 10.1101/2024.08.01.605459

**Authors:** Y. Alonso-Caraballo, Y. Li, N.J. Constantino, M.A. Neal, G.S. Driscoll, Y. Manasian, G.K. Cai, M. Mavrikaki, V.Y. Bolshakov, E.H Chartoff

## Abstract

Opioid use disorder is characterized by compulsive drug seeking and heightened relapse vulnerability following abstinence, a phenomenon known as incubation of craving. Although preclinical data suggest similar behavioral expression of opioid use between sexes, conclusive evidence on sex differences in craving and relapse across abstinence periods remains lacking. Here, we investigated the effects of abstinence from oxycodone self-administration on neurotransmission in the paraventricular thalamus (PVT) to nucleus accumbens shell (NAcSh) pathway in male and female rats. Using optogenetics and ex vivo electrophysiology, we assessed synaptic strength, glutamate release probability, and intrinsic excitability of NAcSh medium spiny neurons (MSNs) following 1 (acute) or 14 (prolonged) days of forced abstinence. No sex differences were observed in oxycodone self-administration or somatic withdrawal. However, females exhibited greater cue-induced relapse after prolonged but not acute abstinence. Prolonged abstinence produced comparable increases in PVT-NAcSh synaptic strength and presynaptic glutamate release probability in both sexes, while inhibitory transmission and MSN excitability were largely unaltered. The dissociation between comparable circuit-level plasticity and sex-specific relapse vulnerability suggests that PVT-NAcSh strengthening represents a shared neuroadaptation to oxycodone abstinence, while mechanisms driving heightened relapse in females likely involve additional circuit elements that remain to be identified.

## Introduction

Relapse is a hallmark of substance use disorder [1,2]. Relapse-inducing triggers include cues previously associated with the drug, stressful events and/or environments, and taking the drug itself (i.e., drug priming). These triggers can result in resumption of drug-taking (relapse), despite negative consequences [3–6]. Several preclinical models of relapse have been developed, and the model most closely mimicking the human condition involves drug self-administration, an operant behavior in which the animal is required to press a lever to obtain the drug and can therefore regulate its drug intake. To specifically assess relapse-like behavior, animals undergo a period of “forced abstinence” after several weeks of drug self-administration. During forced abstinence, the animals are returned to their home cage without access to the drug [7]. When animals are re-introduced to the self-administration behavioral chambers after forced abstinence, they engage in rapid lever pressing, even though no drug is delivered in response. This time-dependent increase in motivation to seek out drug is referred to as ‘incubation of craving’ that leads to relapse [8–12]. Following extended periods of abstinence, incubated craving generally remains elevated before gradually stabilizing and declining[10]. The use of extinction training is also a common approach to study relapse after self-administration. In contrast to forced abstinence, extinction training prompts animals to learn that the drug is no longer available based on their actions (instrumental responding, e.g. lever presses or nose pokes), leading to a gradual reduction in drug-seeking behavior. Both models can trigger relapse when animals are reintroduced to drug-associated cues, but the neural pathways involved and the behavioral outcomes differ [7,13,14]. In these models, extinction training teaches animals that lever pressing no longer results in drug delivery. However, the drug-paired cues (contextual or discrete) are not extinguished. When presented during relapse, the cues elicit increased lever pressing based on the prior drug association formed during self-administration.

Recent studies suggest that glutamatergic projections from the paraventricular nucleus of the thalamus (PVT) to the nucleus accumbens shell (NAcSh) are necessary for the expression of opioid withdrawal signs [15,16] and for cue-induced relapse after abstinence but not extinction [13]. The nucleus accumbens (NAc) is an important node involved in cue-driven reward-seeking behaviors [17–20]. Multiple factors, including synaptic plasticity, contribute to behavioral changes over time. For example, increased synaptic strength in projections from the PVT to D2R-expressing NAc MSNs contributes to naloxone-induced withdrawal behaviors following morphine administration[15], which are evident in the first few days of abstinence and then dissipate. In addition, increased synaptic strength in projections from the PVT to D1R-expressing NAc MSNs is associated with relapse to heroin-seeking after 14 days of abstinence[13]. It has also been found that glutamatergic strength decreases from PVT to parvalbumin interneurons (PV-IN) after self-administration and during extinction and that rescuing this projection prevents heroin relapse [21]. Although it is clear that changes in glutamatergic inputs onto MSNs and other interneurons may contribute to opioid withdrawal and reinstatement of drug-seeking [13,15,22,23], the relationships between NAcSh MSN circuitry functions, length of abstinence from oxycodone self-administration, level of drug-seeking, and sex specificity remain unclear.

We aimed to determine whether synaptic strength in PVT-NAcSh projections is affected following oxycodone abstinence and whether such changes are associated with cue-induced relapse and drug-seeking. Additionally, we examined whether there are sex-specific differences in either cue-induced relapse or PVT-NAcSh synaptic transmission after either 1 (acute) or 14 (prolonged) days of forced abstinence. Our results demonstrate that sex-specific enhancement in cue-induced relapse emerges after prolonged abstinence but not during acute abstinence from oxycodone self-administration. Although both males and females show increased cue-induced relapse after prolonged abstinence, females exhibited a greater relapse rate compared to males. Both sexes showed similar increases in PVT-NAcSh synaptic strength after prolonged abstinence, while synaptic strength was not altered after acute abstinence compared to saline controls. Together, these findings reveal a time-dependent increase in PVT-NAcSh synaptic strength and a sex-specific effect of prolonged abstinence on cue-induced relapse, while synaptic enhancements after prolonged abstinence were not sex-specific.

## Results

### Male and female rats escalate oxycodone intake under long access self-administration

Female (Saline, N=22; Oxycodone N=23) and male (Saline, N=15; Oxycodone, N=27) rats were trained to self-administer oxycodone (or saline) under short access (ShA) conditions (1h/day for 8 days), followed by 14 days of long access (LgA) conditions (6 h/day) (**Fig. 1A Experimental Design**). Males and females escalated their oxycodone intake under LgA conditions: there was an increase in the number of infusions over time (**Fig. 1B**: Mixed-effects model (REML): days x treatment interaction: F_(63,1463)_ = 13.14, p < 0.0001; main effect of days: F_(21,1463)_ = 38.91, p < 0.0001; main effect of treatment: F_(3,71)_ = 31.29, p < 0.0001; no sex difference: F_(1,71)_ = 0.04, p = 0.84), and there was an increase in active lever presses (**Fig. 1C**: Mixed effects model (REML): days x treatment interaction: F_(63,1149)_ = 5.210, p < 0.0001; main effect of days: F_(21,1149)_ = 14.59, p < 0.0001; main effect of treatment: F_(3,56)_ = 12.48, p < 0.0001; no sex differences: F_(1,56)_ = 0.40, p = 0.53). There was no change in inactive lever presses in either sex and lever pressing did not increase across days (**Fig. 1D**: Mixed effects model; main effect of treatment: F_(3,56)_ = 3.909, p = 0.01; no change across days: F_(21,1148)_ = 1.47, p = 0.08; no sex difference: F_(1,56)_ = 0.41, p = 0.52). These findings are consistent with published results [33], showing that escalation of oxycodone self-administration behavior did not differ between male and female rats under LgA conditions.

**Figure 1.**
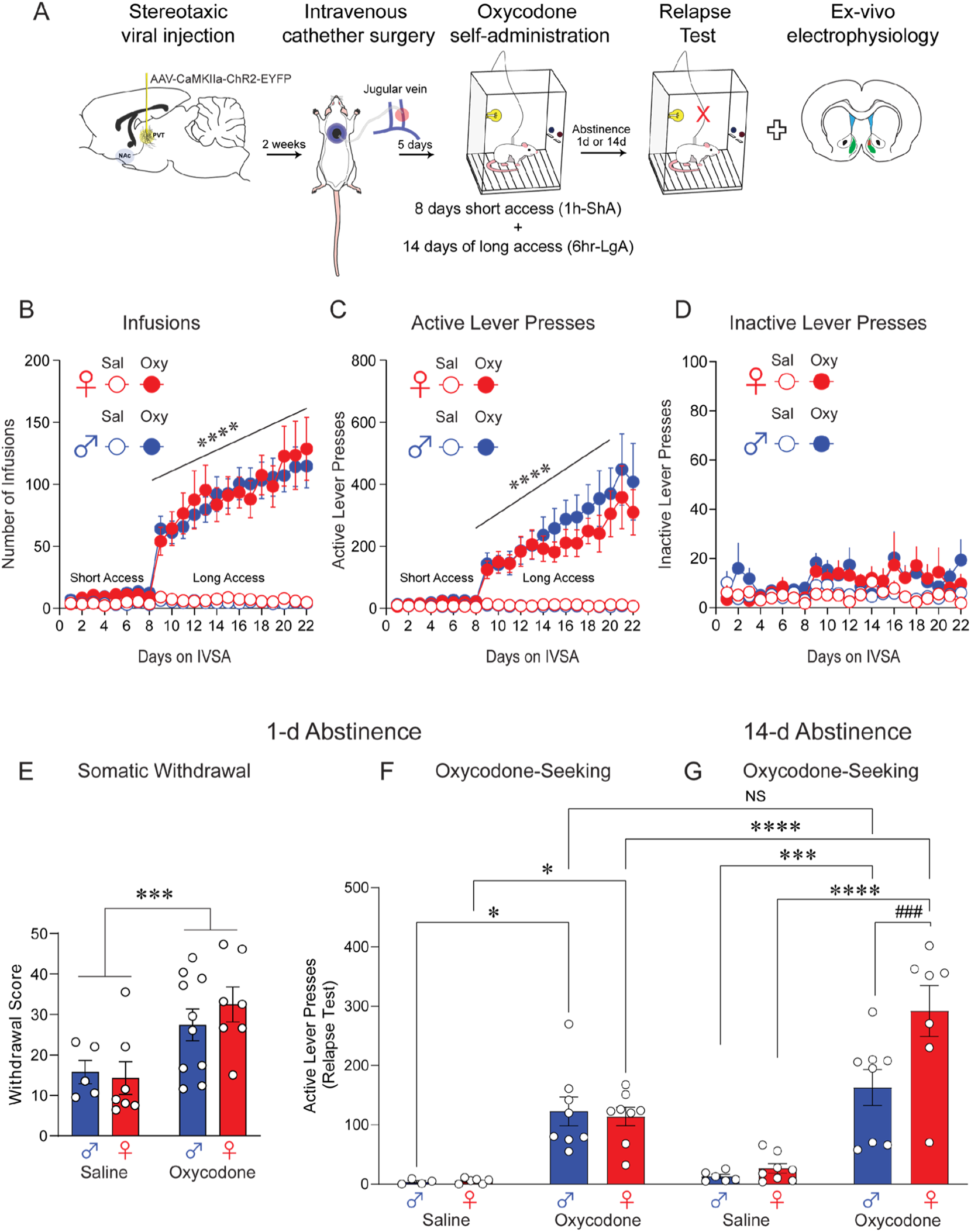
Oxycodone self-administration and oxycodone-seeking after short and prolonged abstinence. **A**: Experimental Timeline. PVT intracranial injections of AAV5-CAMKIIɑ-ChR2-EYFP, followed by intravenous catheter surgery and 22 days of oxycodone self-administration. Rats then received short (24 hours) or long (14 days) abstinence from oxycodone self-administration and tested for cue-induced oxycodone relapse test. Rats were euthanized, and brain slices were prepared for ex-vivo electrophysiological recordings. **B**: Number of infusions across short and long-access oxycodone self-administration in male and female rats. Mixed-effects model: main effect of days (increased infusions over time): p> 0.0001, main of treatment: p < 0.0001, no effect of sex: p = 0.84. **C**: Active lever presses across short- and long-access oxycodone self-administration. Mixed-effects model: main effect of days (increased infusions over time): p> 0.0001, main of treatment: p < 0.0001, no effect of sex: p = 0.53. **D**: Inactive lever presses across short- and long-access oxycodone self-administration. No significant differences between males and females or treatments were observed. **E**: Somatic withdrawal signs increased in both males and females. Two-way ANOVA; main effect of treatment: ***p = 0.002. **F**: Oxycodone-seeking (relapse test) after acute abstinence. Similar oxycodone-seeking behaviors between males and females but increased oxycodone-seeking compared to saline after acute abstinence in males and females. Two-way ANOVA; Sidak’s multiple comparisons females saline vs. oxycodone *p = 0.01; males saline vs. oxycodone *p = 0.02. **G**: Oxycodone-seeking after prolonged abstinence. Both males and females exhibited increased drug-seeking compared to saline. Two-way ANOVA: main effect of drug: F(3,47) = 36.3 *** = p < 0.0001. Females exhibited significantly higher oxycodone-seeking behaviors compared to males: Sidak’s multiple comparisons ### p = 0.006. Data is shown as mean ± SEM.

To assess whether our LgA oxycodone SA produced similar levels of dependence in male and female rats, we measured spontaneous somatic withdrawal 1 day after the last LgA oxycodone SA session in females (n = 7 saline, 7 oxycodone) and males (n = 5 saline, 10 oxycodone). Since opioid dependence is characterized by somatic withdrawal signs upon withdrawal, we predicted an increase in somatic withdrawal signs in both sexes. We found a significant increase in expression of somatic withdrawal behaviors (combined into a Total Withdrawal Score) in females and males that received oxycodone compared to saline **(Fig. 1E**: Two-way ANOVA; main effect of treatment: F_(1, 25)_ = 12.68, p = 0.002). There were no sex differences in the somatic withdrawal score (p = 0.67), consistent with the similar level of escalation of oxycodone SA. This is consistent with previously published studies that have shown increases in somatic withdrawal symptoms after 24-hours in male and female rats [34].

### Acute abstinence (1-day) from oxycodone self-administration increased cue-induced oxycodone-seeking in female and male rats

We also assessed cue-induced drug-seeking after acute abstinence from LgA oxycodone SA. Female (n = 6 saline, 8 oxy) and male rats (n = 4 saline, 8 oxy) remained in their home cages without any access to oxycodone (forced abstinence) for 1-day. After 1-day of abstinence, rats returned to the operant box where only the house-cue, lever-cue and lever were available for 2 hrs. No infusions were given. We measured the number of active lever presses during cue presentation without drug delivery. We found that 1-day of abstinence increased cue-induced oxycodone seeking in males and females compared to saline (**Fig. 1F**: Two-way ANOVA; significant sex x treatment interaction: F_(3,47)_ = 3.9, p = 0.02; no effect of sex: F_(1,47)_ = 3.9, p =0.06; main effect of drug: F_(3,47)_ = 36.3, p < 0.0001). Sidak’s multiple comparison: female saline group vs. oxycodone group, p = 0.01; male saline group vs. oxycodone group, p = 0.02.

### Cue-induced drug-seeking after prolonged abstinence (14-days) exhibited sex-specific differences

Cue-induced drug-seeking was assessed after 14-days of abstinence from LgA oxycodone SA. Female (n = 8 saline, 7 oxycodone) and male (n = 6 saline, 8 oxycodone) rats underwent 14-days of forced abstinence in their home cages. On the 14th day of abstinence, rats were returned to the operant box where only the house-cue, lever-cue and lever were available for 2 hrs. No infusions were given. We found that both male and female rats on oxycodone pressed the active lever more frequently than rats in the saline control group (**Fig. 1G**: Two-way ANOVA: main effect of drug: F (3,47) = 36.3, p < 0.0001. Additionally, females made significantly more active lever presses than males (**Fig. 1G**: Two-way ANOVA: significant drug x sex interaction: F (3,47) = 3.9, p = 0.02; Sidak’s multiple comparisons test female oxycodone greater than male oxycodone, p = 0.006). In addition, when comparing oxycodone-seeking in males and females during acute or prolonged abstinence, we found that females but not males show an increase in seeking over time (**Fig. 1F-G**: Two-way ANOVA: significant day x sex interaction: F (3, 47) = 3.9, p = 0.02. Sidak’s multiple comparisons: oxycodone male acute vs. prolonged: p = 0.9; oxycodone female acute vs. prolonged: p < 0.0001.

These findings suggest that prolonged forced abstinence increased oxycodone-seeking in both males and females, with a stronger effect in females than in males.

### Electrophysiological characterization of PVT-NAcSh projections

To test whether PVT-arising inputs to NAcSh contribute to cue-induced relapse and drug-seeking, we targeted PVT-NAcSh projections optogenetically by expressing ChR2 in the PVT via stereotaxic injections of AAV5-CaMKIIα-ChR2-eYFP viral construct in rats from all experimental groups. 7-8 weeks after viral transfection, eYFP-tagged ChR2 was expressed at the injection site (PVT) and ChR2-eYFP-expressing PVT-arising projections were found in the NAcSh (**Fig. 2A**). Using whole-cell patch clamp recordings (**Fig. 2B**), we confirmed that PVT projections form functional synapses on the NAcSh neurons (**Fig. 2C, D**). Photostimulation-induced (470 nm. 5 ms-long pulses of blue light) excitatory postsynaptic currents (EPSCs) were recorded under voltage clamp recording conditions at a holding potential of –70 mV. EPSCs at the PVT-NAcSh synapses are glutamatergic, as demonstrated by their sensitivity to NBQX (10µM), an AMPA/Kainate receptor antagonist, and D-AP5, an NMDA receptor antagonist (**Fig. 2D, E**). Glutamatergic EPSCs in PVT-NAcSh projections were monosynaptic in nature as they were rescued by the potassium channel blocker, 4-AP (1 mM) after they were blocked by tetrodotoxin (TTX, 1 µM), a sodium channel blocker (**Fig. 2G, H**) [35].

**Figure 2.**
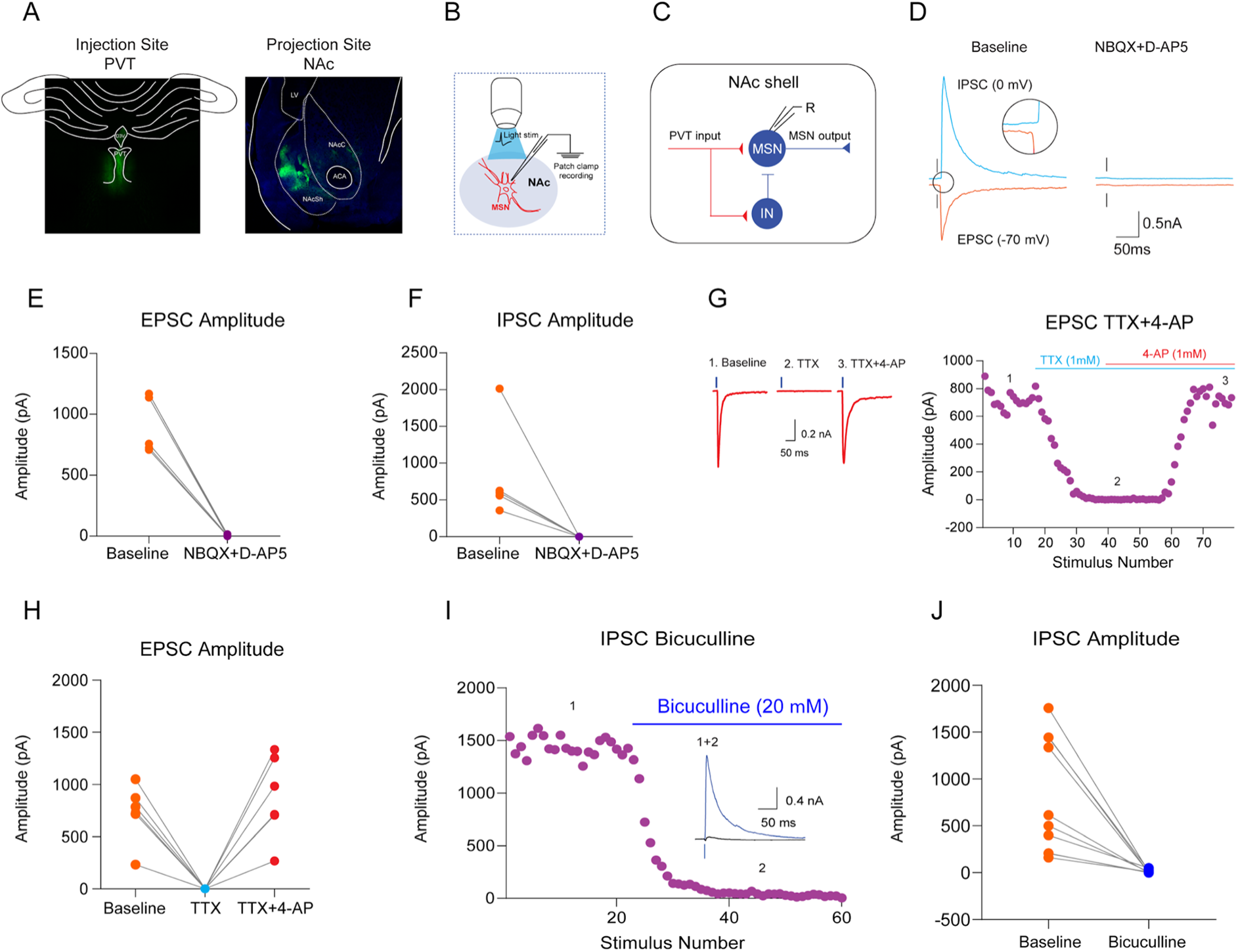
Synaptic Properties in PVT-to-NAcSh Projections. **A**: Representative images showing expression of ChR2-EYFP at the injection site in PVT and ChR2-EYFP-expressing projecting fibers in NAcSh. **B**: Schematic of optical stimulation of PVT terminals in NAcSh and recording of optically-evoked EPSCs in MSNs. **C**: Schematic representation of the local circuit within NAcSh with PVT afferents forming monosynaptic glutamatergic contacts on both MSNs and GABAergic interneurons, resulting in of feedforward inhibitory responses in MSNs. **D:** Photostimulation-induced (10.5 mW/mm^2^; 5 ms) EPSC (orange) and IPSC (cyan) recorded from NAcSh MSN at holding potentials of -70 mV or 0 mV, respectively. Recordings were performed under control conditions first (left; baseline) and 10 min after NBQX (10 μM) and D-AP5 (50 μM; right) were added to the bath solution. The inset shows a delayed onset (synaptic latency) of the IPSC recorded at 0 mV. **E-F:** Summary plot of the amplitude of EPSC (panel E) and IPSC (panel F) recorded in MSNs under control conditions (baseline) and after bath application of NBQX and D-AP5. The symbols represent individual experiments. **G:** Rescue of light-induced and TTX-blocked EPSCs at PVT-NAcSh projections by 4-AP. Left, an example of recordings shows EPSC (average of 10 traces) recorded at -70 mV under control conditions (baseline; 1), the EPSC was blocked by TTX (1 μM, 2), and application of 4-AP (1 mM) in the continuing presence of TTX restored the EPSC (3), thus confirming the monosynaptic nature of the PVT-NAcSh projections. Right, the time course of the EPSC amplitude changes. **H:** Summary plot of the experiments showing the EPSC amplitudes in NAcSh MSNs under three conditions (baseline, TTX, and TTX + 4-AP. **I:** Feed-forward IPSCs in NAcSh MSNs, recorded at 0 mV, were blocked by the GABA_A_ receptor antagonist bicuculline (20 μM). **J:** IPSC amplitudes recorded from NAcSh under control conditions (baseline) and after application of bicuculline.

Stimulation of glutamatergic PVT projections resulted in activation of GABAergic interneurons in NAcSh **(Fig. 2C)**, triggering feed-forward inhibitory postsynaptic currents (IPSCs) in recorded MSNs **(Fig. 2D).** The IPSCs, recorded at a holding potential of 0 mV, were GABAergic as they were blocked by the GABA_A_ receptor antagonist, bicuculline (20 μM) (**Fig. 2I, J**), and they were disynaptic in nature (**Fig. 2C**) as they were sensitive to glutamate receptors antagonists NBQX and D-AP5 (**Fig. 2F**). Based on these findings, we conclude that PVT sends monosynaptic projections to NAcSh, forming glutamatergic synapses on MSNs, as well as triggering feedforward inhibition in the PVT-NAcSh circuits.

### Synaptic strength in PVT-NAcSh projection is increased after prolonged but not acute abstinence in both male and female rats

To examine the effect of acute abstinence from oxycodone self-administration on synaptic transmission in the PVT-NAcSh projections, we performed whole-cell patch-clamp recordings of light-induced EPSCs in medium spiny neurons in NAcSh in slices from rats which self-administered oxycodone at both abstinence time points. To assay the effects of acute abstinence on the efficacy of excitatory synaptic transmission, we obtained input-output curves for the light-induced EPSCs in all experimental groups (male saline group: 5 rats, 18 cells; male oxycodone group: 8 rats, 36 cells; female saline group: 7 rats; 26 cells; female oxycodone group: 6 rats, 20 cells) by recoding the EPSCs at a holding potential of –80 mV evoked by photostimuli of increasing intensity (**Fig. 3A**). Although the EPSC amplitude increased with light density (**Fig. 3A**: Three-way ANOVA: main effect of light intensity F_(5,582)_ = 52.69, p < 0.0001), we did not find any treatment effect on synaptic efficacy in the PVT-NAcSh pathway (**Fig. 3A**: Three-way ANOVA: no effect of treatment: F_(1,582)_ = 1.5, p = 0.2) or sex specificity (**Fig. 3A**: Three-way ANOVA: no effect of sex: F_(1,582)_ = 0.5, p = 0.5). This finding indicates that 1-day abstinence from oxycodone self-administration did not affect the efficacy of glutamatergic synaptic transmission in PVT-NAcSh projections in both male and female rats.

**Figure 3.**
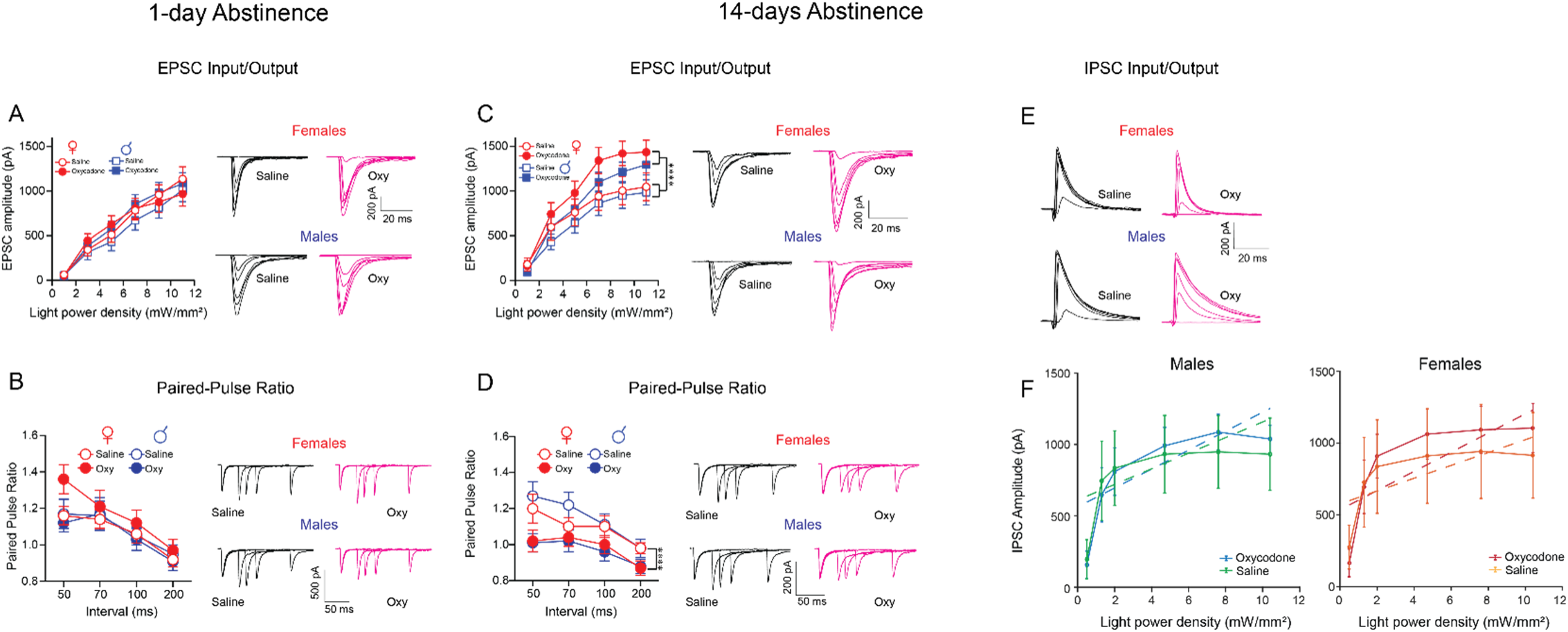
Prolonged abstinence from oxycodone self-administration is associated with increased synaptic strength in glutamatergic PVT projections to MSNs in NAcSh in male and female rats. **A: Right**, 1-day of abstinence from oxycodone self-administration (acute abstinence) had no effect on the efficacy of glutamatergic synaptic transmission in PVT projections to MSNs in NAcSh, as assessed with synaptic input-output curves for light-induced EPSCs which were triggered by the pulses of blue light of increasing intensity. There was no significant difference between saline (n = male: 5 rats, 18 cells; females: 7 rats, 26 cells) and oxycodone (n = male: 8 rats, 36 cells; females: 6 rats, 20 cells) groups. Three-way ANOVA no effect of treatment: p = 0.2, no effect of sex: p = 0.5. **Right**, example traces of EPSCs triggered by light pulses of increasing intensity from different experimental groups. **B: Left**, the magnitude of the paired-pulse ratio remained unchanged in the oxycodone (n = males: 10 rats, 25 cells; females: 6 rats, 22 cells) compared to saline control (n = males: 5 rats, 28 cells; females: 7 rats, 30 cells) groups, indicating that the probability of glutamate release was not affected after acute abstinence from oxycodone self-administration. Three-way ANOVA no effect of drug, p = 0.3, no effect of sex, p = 0.1. **Right**, representative traces of EPSCs evoked by paired-pulses of blue light (10.5 mW/mm²) at different interpulse intervals (50, 70, 100 and 200 ms). **C: Left**, the efficacy of glutamatergic synaptic transmission in PVT projections to MSNs in NAcSh, assessed as in **A** with synaptic input-output curves, was enhanced in both male and female rats in the oxycodone group (n = males: 7 rats, 20 cells; females: 5 rats, 21 cells) compared to saline control group (n = males: 5 rats, 16 cells; females: 7 rats, 21 cells) after 14-days of abstinence from oxycodone self-administration (prolonged abstinence). Three-way ANOVA main effect of drug ****p < 0.0001. **Right**, example traces of EPSCs from different experimental groups. **D: Left**, the magnitude of the paired-pulse ratio decreases in the oxycodone groups (n = males: 7 rats, 33 cells; females: 6 rats, 22 cells) compared to saline control groups (n = males: 5 rats, 23 cells; females: 7 rats, 23 cells). Three-way ANOVA, main effect of drug **** p < 0.0001. **E:** Example traces of IPSCs from different experimental groups triggered by photostimuli of increasing intensity. The IPSCs were recorded at a holding potential of 0 mV. **F: Left (males),** linear mixed-effects modeling revealed a significant main effect of light intensity on IPSC amplitude (F_(1, 158)_ = 43.49, p = 6.1 × 10⁻¹⁰), indicating robust recruitment of inhibitory synaptic responses with increasing stimulation. No significant main effect of condition was observed in males (F_(1, 158)_ = 0.00023, p = 0.988), and no Light × Condition interaction was detected (F_(1, 158)_ = 0.543, p = 0.462), indicating that prolonged abstinence from oxycodone does not alter baseline inhibitory strength or recruitment dynamics in male NAcSh MSNs. **Right (females)**, linear mixed-effects modeling revealed a significant main effect of light intensity on IPSC amplitude (F_(1, 164)_ = 21.26, p = 8.0 × 10⁻⁶), indicating robust recruitment of inhibitory synaptic responses with increasing stimulation. No significant main effect of condition was observed in females (F_(1, 164)_ = 0.025, p = 0.875), and no Light × Condition interaction was detected (F_(1, 164)_ = 1.298, p = 0.256), indicating that prolonged abstinence from oxycodone does not alter baseline inhibitory strength or recruitment dynamics in female NAcSh MSNs. Together, these data indicate that the efficacy of feed-forward inhibition in the PVT–NAcSh pathway, as assessed by input-output curves for light-induced IPSCs, is unaffected by prolonged oxycodone abstinence in either sex. Saline (n = males: 5 rats, 11 cells; females: 7 rats, 14 cells) and oxycodone (n = males: 7 rats, 16 cells; females: 5 rats, 14 cells).

Consistent with the lack of changes in synaptic strength, the magnitude of paired-pulse ratio (PPR), and index of presynaptic function [35] of EPSC amplitude at the PVT-NAcSh synapses did not differ between the treatment groups (male saline: 5 rats, 28 cells; male oxycodone: 10 rats, 25 cells; female saline: 7 rats, 30 cells; female oxycodone: 6 rats, 22 cells) (**Fig. 3B**: Three-way ANOVA: F_(1,443)_ = 1.19, p = 0.3) or sex specificity (**Fig. 3B**: Three-way ANOVA: F_(1,443)_ = 2.20, p = 0.1), indicating that the probability of neurotransmitter release at the PVT-NAcSh synapses was unaffected by acute abstinence.

In analogously designed experiments, we assayed the effects of prolonged abstinence on synaptic transmission in PVT-NAcSh projections. As above, input-output curves for PVT-NAcSh EPSCs were obtained by recording photostimulation-induced EPSCs in MSNs at a holding potential of –80 mV **(Fig. 3C)** (male saline group: 5 rats, 16 cells; male oxycodone group: 7 rats, 20 cells; female saline group: 7 rats; 21 cells; female oxycodone group: 5 rats, 21 cells). We found that the EPSC amplitude increased with light intensity (**Fig. 3C**: Three-way ANOVA: main effect of light intensity; F_(5,442)_ = 40.5, p < 0.0001), and, unlike 1-day abstinence, long abstinence from oxycodone self-administration was associated with increases in the EPSC amplitude in both male and female rats compared to control group (**Fig. 3C**: Three-Way ANOVA: main effect of treatment; F_(1,442)_ = 18.3, p < 0.0001). The effects of prolonged abstinence on synaptic strength in the PVT-NAcSh pathway was similar between males and females.

Notably, the magnitude of PPR at the studied synapses was found to be decreased after prolonged abstinence from oxycodone self-administration in both male and female rats (**Fig. 3D**: Three-Way ANOVA: main effect of treatment: F_(1,388)_ = 25.8, p < 0.001; male saline: 5 rats, 23 cells; male oxycodone: 7 rats, 33 cells; female saline: 7 rats, 23 cells; female oxycodone: 6 rats, 22 cells). Thus, the observed synaptic potentiation in PVT-NAcSh projections after longer abstinence periods was at least in part due to increased glutamate release probability.

To assess whether prolonged abstinence from oxycodone self-administration alters feed-forward inhibition within the PVT-NAcSh pathway (**Fig. 3E-F**), light-evoked IPSC amplitudes were analyzed using linear mixed-effects modeling. In both males and females, IPSC amplitude increased robustly with light intensity, indicating preserved recruitment of inhibitory synaptic responses with increasing stimulation (males: F_(1, 158)_ = 43.49, p = 6.1 × 10⁻¹⁰; females: F_(1, 164)_ = 21.26, p = 8.0 × 10⁻⁶). No significant main effect of treatment condition was detected in either sex (males: F_(1, 158)_ = 0.00023, p = 0.988; females: F_(1, 164)_ = 0.025, p = 0.875), indicating that prolonged abstinence from oxycodone did not alter baseline inhibitory synaptic strength within the PVT-NAcSh pathway in males or females. No significant Light × Condition interactions were observed in either sex (males: F_(1, 158)_ = 0.543, p = 0.462; females: F_(1, 164)_ = 1.298, p = 0.256), indicating that the scaling of inhibitory responses with increasing stimulation intensity was not altered by oxycodone exposure in either sex. Together, these results demonstrate that feed-forward inhibition within the PVT-NAcSh pathway is preserved following prolonged abstinence from oxycodone, with no significant changes in either the magnitude or the input-output scaling of inhibitory synaptic responses in males or females.

Taken together with the observed strengthening of excitatory synaptic transmission in PVT–NAcSh projections after prolonged abstinence (**Fig. 3C**), these results suggest that prolonged abstinence from oxycodone selectively potentiates excitatory but not inhibitory synaptic transmission within this pathway. This relative shift in the excitation–inhibition balance may facilitate MSN output within relapse-related circuitry and contribute to the behavioral effects observed following prolonged abstinence.

### Abstinence from oxycodone self-administration does not change AMPAR subunit composition or AMPAR/NMDAR EPSC amplitude ratio in glutamatergic PVT projections to MSNs in NAcSh

To explore whether changes in glutamatergic PVT projections to MSNs in NAcSh correlated with abstinence from oxycodone self-administration, we recorded light-induced AMPA receptor-mediated (AMPAR) EPSCs at the PVT-NAcSh synapses in slices from all experimental groups at holding potentials of −70 mV, 0 mV or +40 mV. In these experiments, we included endogenous polyamine spermine (200 μM), in the internal recording solution. We then calculated the rectification index for AMPAR EPSCs at the studied PVT-NAcSh projections [36] by dividing the peak amplitude of AMPAR EPSC at +40 mV by the EPSC amplitude at -70 mV. The changes in rectification index associated with experimental interventions would indicate that the AMPAR subunit composition was modified, as the GluR1 subunit trafficking to synapses was shown to affect the rectification index [36]. However, after acute abstinence, we did not observe significant changes in the rectification index (male saline: 5 rats, 23 cells; male oxycodone: 8 rats, 28 cells; female saline: 7 rats, 14 cells; female oxycodone: 6 rats, 18 cells) by sex (**Fig. 4A**, inset bar graph: Two-way ANOVA: F_(1,68)_ = 0.83, p = 0.4) or treatment (**Fig.4A**, inset bar graphs: Two-way ANOVA: F_(1,68)_ = 0.35, p = 0.6).

**Figure 4.**
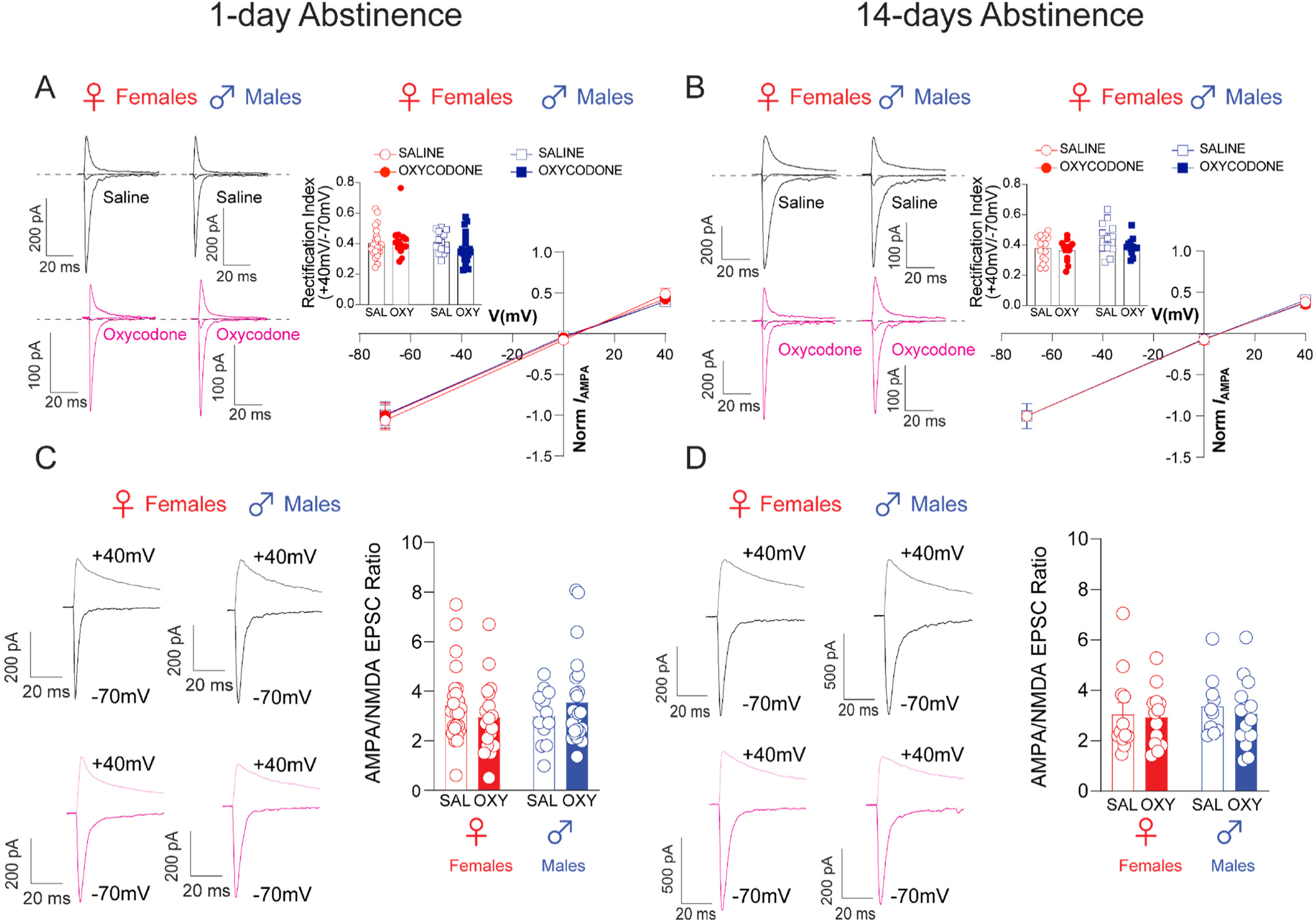
Abstinence from oxycodone self-administration does not change AMPAR subunit composition or AMPAR/NMDAR EPSC amplitude ratio in glutamatergic PVT projections to MSNs in NAcSh. A-B: Rectification index and current-voltage relationship for AMPAR EPSCs for short (A; 1-day) and long (B; 14-days) abstinence periods. **A: Left**, representative traces of AMPAR EPSCs recorded at holding potentials of -70, 0 and +40 mV during acute abstinence. **Right**, current/voltage relationship of AMPAR EPSCs recorded at holding potentials of -70, 0 and +40 mV of saline (n = males: 5 rats, 23 cells; females: 7 rats, 14 cells) vs. oxycodone (n = males: 8 rats, 28 cells; females: 6 rats, 18 cells) rats. **Inset:** Rectification index for EPSCs (calculated as the ratio of peak EPSC amplitudes at +40/-70 mV [EPSC_+40_/EPSC_-70_]). **B: Left**, representative traces of AMPAR EPSCs recorded at holding potentials of -70, 0 and +40 mV during prolonged abstinence. **Right**, current/voltage relationship of AMPAR EPSCs recorded at holding potentials of -70, 0 and +40mV of saline (n = males: 4 rats, 15 cells; females: 5 rats, 22 cells) vs. oxycodone (n = males: 5 rats, 23 cells; females: 5 rats, 15 cells) rats. The recordings were performed in the presence of the NMDA receptor antagonist D-APV (50 μM) in the external medium and spermine (200 μM) in the pipette solution. There were no significant differences between groups. Data is shown as mean ± SEM. Comparisons were made using two-way ANOVAs p > 0.05. **C-D:** AMPA/NMDA ratios of AMPAR EPSCs for short (C) and long (D) abstinence periods. **C: Left**, representative traces of light-induced EPSCs in projections from PVT to MSNs in NAcSh at +40mV (light-colored traces) and -70mV (dark-colored traces) in slices from male and female rats during acute abstinence. **Right**, AMPAR/NMDAR EPSC amplitude ratios for saline (n = males: 5 rats, 14 cells; females: 7 rats, 24 cells) and oxycodone (males: 8 rats, 24 cells; females: 6 rats, 18 cells) groups during acute abstinence. **D: Left**, representative traces of light-induced EPSCs in projections from PVT to MSNs in NAcSh at +40mV (light-colored traces) and -70mV (dark-colored traces) in slices from male and female rats during prolonged abstinence. **Right**, AMPAR/NMDAR EPSC amplitude ratios for saline (n = males: 5 rats, 12 cells; females: 5 rats, 15 cells) and oxycodone (males: 5 rats, 12 cells; females: 5 rats, 15 cells) groups. Data is shown as mean ± SEM. Comparisons were made using two-way ANOVA.

Similarly, after prolonged abstinence we did not observe significant changes in the rectification index (male saline: 4 rats, 15 cells; male oxycodone: 5 rats, 23 cells; female saline: 5 rats, 22 cells; female oxycodone: 5 rats, 15 cells) by treatment (**Fig. 4B** (inset bar graph): Two-way ANOVA: F_(1,50)_ = 2.7, p = 0.1). These results suggest that neither acute nor prolonged abstinence from oxycodone self-administration was associated with changes in the AMPAR subunit composition at synapses formed by PVT projecting fibers on MSNs in NAcSh. Thus, synaptic potentiation in PVT-NAcSh projections, observed after 14 days of abstinence (see Fig. 3C), was unlikely due to increased GluR1 trafficking at the synapses studied.

Consistent with the latter notion, the AMPAR/NMDAR EPSC amplitude ratio was not affected by acute abstinence (**Fig. 4C**: Two-way ANOVA: no effect of sex: F_(1,76)_ = 0.06, p = 0.8; no effect of treatment: F_(1,76)_ = 0.008, p = 0.9; n = male saline: 5 rats, 14 cells; male oxycodone: 8 rats, 24 cells; female saline: 7 rats, 24 cells; female oxycodone: 6 rats, 18 cells). The AMPAR/NMDAR EPSC ratio was assessed by measuring the amplitude of the NMDAR-mediated EPSC at a holding potential of +40mV, 40ms after the peak of AMPAR-mediated EPSC at -70mV. Similarly, the AMPAR/NMDAR EPSC amplitude ratio was also unaffected after prolonged (14-days) abstinence (**Fig. 4D**: Two-way ANOVA: no effect of sex: F_(1,45)_ = 0.4, p = 0.5; no effect of treatment: F_(1,45)_ = 0.3, p = 0.6; (n = male saline: 5 rats, 12 cells; male oxycodone: 5 rats, 12 cells; female saline: 5 rats, 15 cells; female oxycodone: 5 rats, 15 cells), suggesting that acute or prolonged abstinence from oxycodone self-administration might have no detectable postsynaptic effects as assessed by these synaptic measures in PVT-NAcSh projections.

### Prolonged but not acute abstinence from oxycodone self-administration shows a trend toward increased NAcSh MSN intrinsic excitability in males but not females

To determine whether abstinence from oxycodone self-administration alters intrinsic excitability of NAcSh medium spiny neurons (MSNs), we performed whole-cell current-clamp recordings from NAcSh MSNs in slices from saline- (males: 7 rats, 16 cells; females: 4 rats, 7 cells) and oxycodone-treated rats (males: 6 rats, 18 cells; females: 8 rats, 15 cells) following acute abstinence (1 day; **Fig. 5A, C–E**), as well as from saline- (males: 3 rats, 8 cells; females: 6 rats, 14 cells) and oxycodone-treated rats (males: 4 rats, 14 cells; females: 6 rats, 15 cells) following prolonged abstinence (14 days; **Fig. 5B, F–H**). Spike output was quantified as the number of action potentials fired during depolarizing current steps. Across all conditions, injected current robustly increased spike output, confirming intact spike count–current relationships in all groups (acute abstinence males: F_(1, 370)_ = 50.03, p = 7.6 × 10⁻¹²; acute abstinence females: F_(1, 230)_ = 71.63, p = 3.1 × 10⁻¹⁵; prolonged abstinence males: F_(1, 225)_ = 19.23, p = 1.8 × 10⁻⁵; prolonged abstinence females: F_(1, 306)_ = 30.14, p = 8.4 × 10⁻⁸).

**Figure 5.**
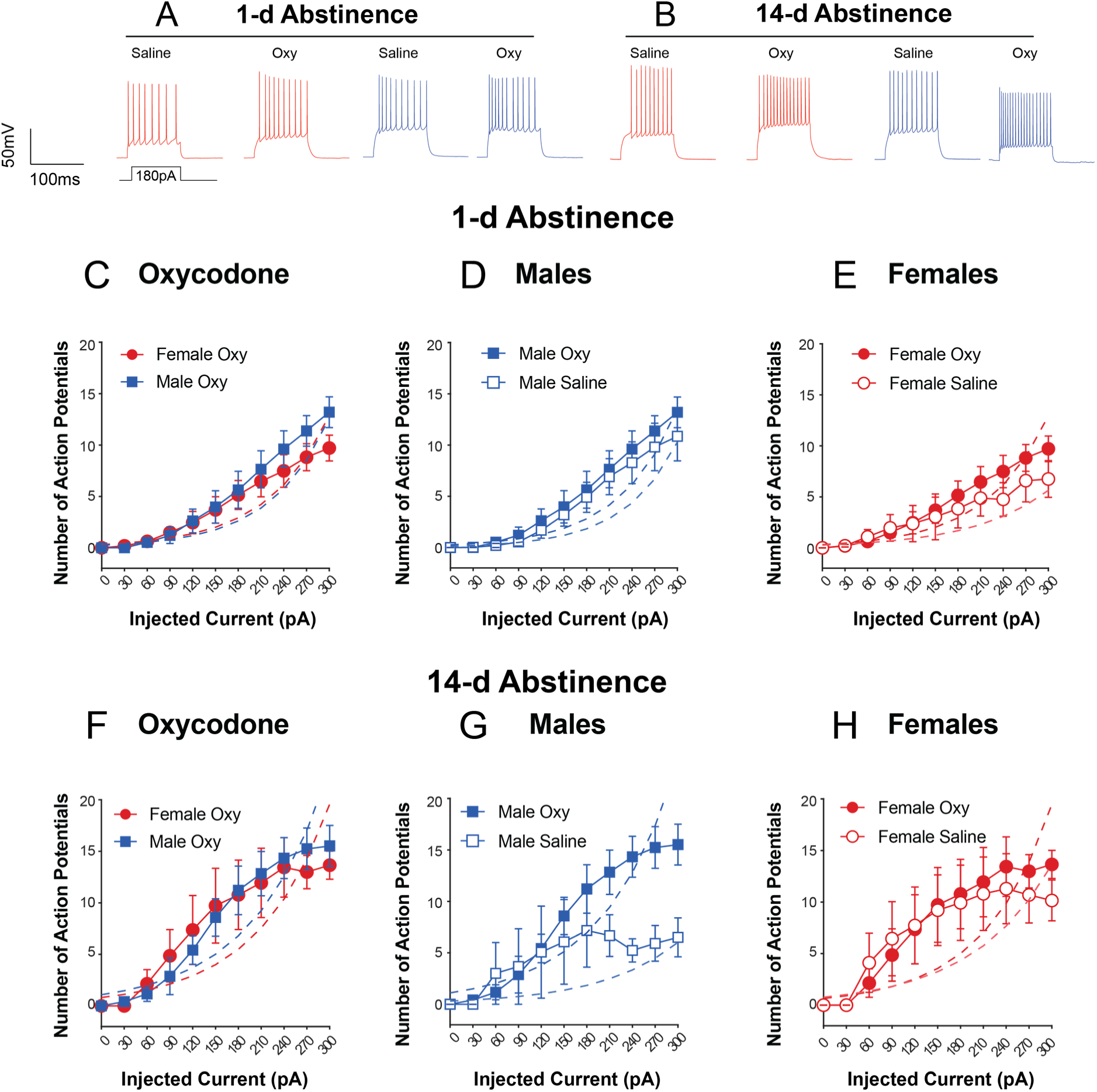
Intrinsic excitability of NAcSh medium spiny neurons is largely unaltered following oxycodone abstinence, with a trend toward increased spike output in males after prolonged abstinence. **A–B**: Example current-clamp traces from NAcSh MSNs from saline- and oxycodone-treated rats after 1 day (**A**) and 14 days (**B**) of abstinence. Sample sizes: 1-day saline (males: 7 rats, 16 cells; females: 4 rats, 7 cells), 1-day oxycodone (males: 6 rats, 18 cells; females: 8 rats, 15 cells), 14-day saline (males: 3 rats, 8 cells; females: 6 rats, 14 cells), 14-day oxycodone (males: 4 rats, 14 cells; females: 6 rats, 15 cells). **C, F**: Spike count–current relationships in oxycodone-treated animals revealed no significant sex differences in baseline spike output or spike output gain at 1 day (**C**: condition F_(1, 351)_ = 0.063, p = 0.802; interaction F_(1, 351)_ = 0.105, p = 0.746) or 14 days of abstinence (**F:** condition F_(1, 300)_ = 0.176, p = 0.675; interaction F_(1, 300)_ = 0.036, p = 0.849). **D–E**: No significant effect of oxycodone on spike output was detected in males or females after 1 day of abstinence (males: condition F_(1, 370)_ = 0.454, p = 0.501; females: condition F_(1, 230)_ = 0.571, p = 0.451). **G:** After 14 days of abstinence, oxycodone-treated males showed numerically higher spike output than saline controls (7.46 vs. 4.43 spikes/step; Cohen’s d = 1.18), but this did not reach statistical significance (condition: F_(1, 225)_ = 2.419, p = 0.121; interaction: F_(1, 225)_ = 0.008, p = 0.930); post-hoc power analysis indicated only ∼20% power with the current sample, and 13 rats per group would be required for 80% power. **H**: No significant effect of oxycodone on spike output was detected in females after 14 days of abstinence (condition: F_(1, 306)_ = 0.025, p = 0.875; interaction: F_(1, 306)_ = 0.366, p = 0.546). Spike count data were analyzed using nested hierarchical Poisson GLMMs fitted separately for each sex and abstinence duration. Data are shown as mean ± SEM.

Following acute abstinence, oxycodone exposure did not significantly alter baseline spike output or spike output gain in males (**Fig. 5D**: condition, F_(1, 370)_ = 0.454, p = 0.501; current × condition, F_(1, 370)_ = 0.236, p = 0.628) or females (**Fig. 5E**: condition, F_(1, 230)_ = 0.571, p = 0.451; current × condition, F_(1, 230)_ = 0.776, p = 0.379), indicating that acute oxycodone abstinence does not detectably alter MSN intrinsic excitability in either sex.

Following prolonged abstinence, no significant effect of oxycodone exposure on baseline spike output or spike output gain was detected in females (**Fig. 5H**: condition, F_(1, 306)_ = 0.025, p = 0.875; current × condition, F_(1, 306)_ = 0.366, p = 0.546). In males, oxycodone-treated animals showed numerically higher spike output compared to saline controls (mean spikes per step: oxycodone 7.46 ± SD 2.29 vs saline 4.43 ± SD 2.82), corresponding to a large effect size (Cohen’s d = 1.18); however, this difference did not reach statistical significance (**Fig. 5G**: condition, F_(1, 225)_ = 2.419, p = 0.121; current × condition, F_(1, 225)_ = 0.008, p = 0.930). A post-hoc power analysis indicated that the current sample size (3 saline rats, 4 oxycodone rats) provided only approximately 20% power to detect this effect, and that 13 rats per group would be required to achieve 80% power. These results should therefore be interpreted with caution, and the possibility of a biologically meaningful increase in MSN excitability in males following prolonged oxycodone abstinence cannot be excluded.

No significant difference in baseline spike output or spike output gain was observed between males and females at either 1 day (**Fig. 5C**: condition, F_(1, 351)_ = 0.063, p = 0.802; current × condition, F_(1, 351_) = 0.105, p = 0.746) or 14 days of abstinence (**Fig. 5F**: condition, F_(1, 300)_ = 0.176, p = 0.675; current × condition, F_(1, 300)_ = 0.036, p = 0.849).

Together, these results indicate that intrinsic excitability of NAcSh MSNs is largely unaltered following oxycodone abstinence in both sexes. The exception is a trend toward increased spike output in males following prolonged abstinence, which, despite a large observed effect size, did not reach statistical significance due to limited sample size in this group.

### Dendritic morphology of NAcSh-MSNs remained unchanged regardless of abstinence duration from oxycodone self-administration

More than 85% of Neurobiotin-stained cells obtained from the medial division of NAcSh in male or female rats are medium spiny neurons (MSNs) characterized by dense spines on a second or higher order of dendritic branches (**Figure 6A-C**). The number of MSNs primary dendrites (3.3-3.6 on average for male rats, 2.8-4.1 in average for female rats) and elongated dendritic trees (∼1.5 on average ratios of long axis over short axis of dendritic field among treatment groups) are consistent with the previously described morphological features of NAcSh-MSNs [32]. Axons arising from soma or proximal dendrites can be observed in the majority of reconstructed MSNs (86.8% or 85.7% in male or female rats, respectively). Some of the axons (21% or 37% in male or female rats) were found to give rise to axonal collateral ramifications with terminal-like varicosities in the vicinity of their soma. This observation suggests that some MSNs may contribute to feedforward inhibition in studied projections as manifested by polysynaptic IPSC activated by photostimulation of PVT afferents (**Figure 6A**, left and **Figure 2C,D,F,I and J** [37,38]). In addition, less than 15% of Neurobiotin-stained neurons were identified either as sparsely spined neurons (SSNs) characterized by a few spines on thread-like dendrites, or aspiny neurons (ASNs) without visible dendritic spine (**Figure 6A-C**). These two types of neurons observed in both female and male rats were not included in the subsequent quantitative analysis.

**Figure 6.**
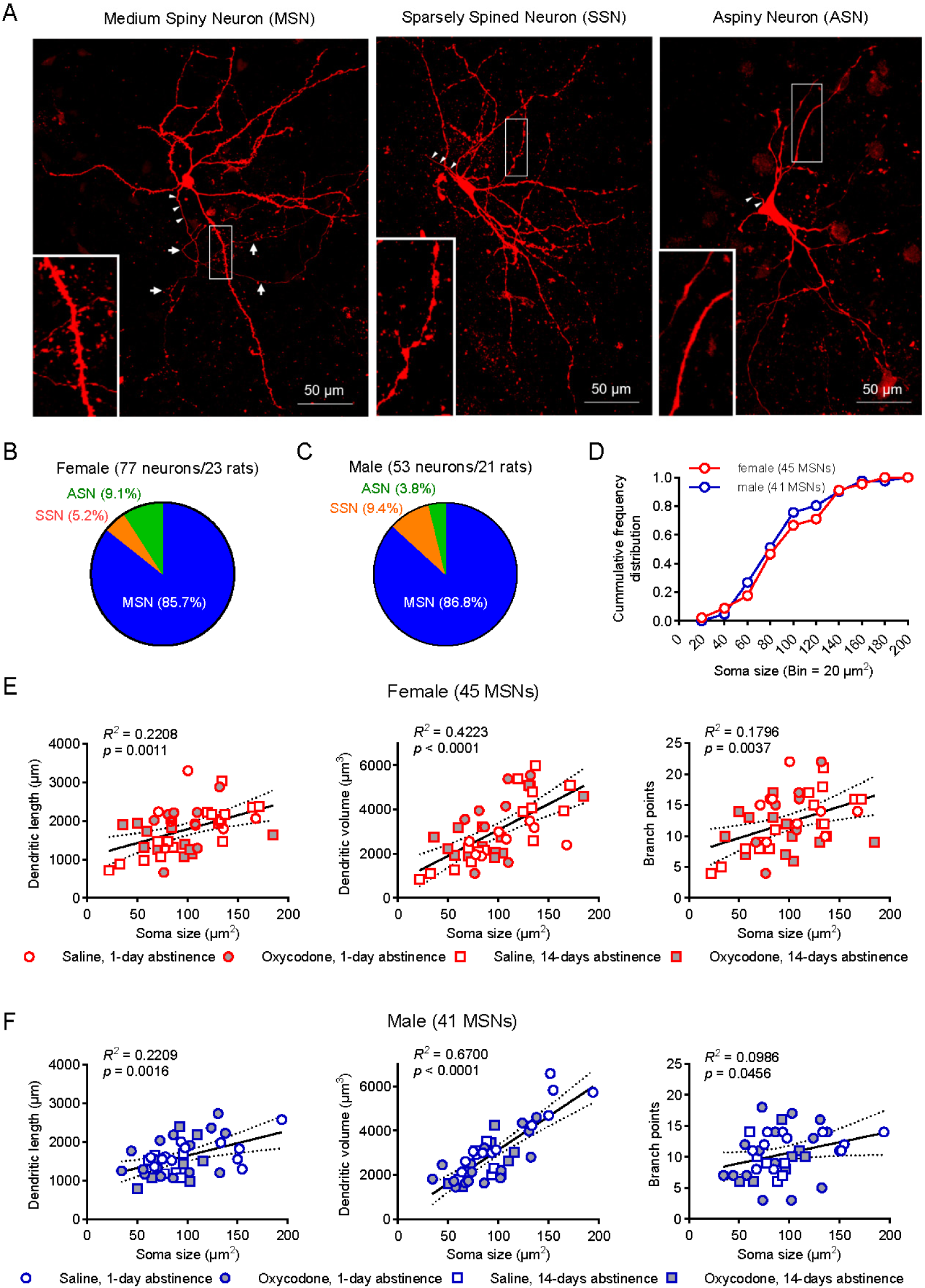
Cell types in the NAcSh and variation in morphology of MSNs. A: Stacked confocal images show examples of a medium spiny neuron (**left**), a sparsely spine neuron (**middle**) and an aspiny neuron (**right**), respectively. Inset images show enlarged view of a segment of dendritic branch and its spines from the same neuron. Arrowheads in three panels indicate the initial segment of axons. Arrows in the left panel indicate the axonal collaterals in the vicinity of its soma. **B-C**: Summary plot shows the percentages of three types of neurons observed in NAcSh in female and male rats. D: There is no difference in cumulative frequency distribution of soma size of examined MSNs between female (**red circles**) and male (**blue circles**) rats (p = 0.954, Mann-Whitney test). **E-F**: Correlation analysis of soma sizes versus their dendritic length (**left**), dendritic volume (**middle**), or branch points (**right**) in female (**E**) and male rats (**F**). The values of the correlation coefficient (R2) and significance (p) are shown in each panel. The solid line in each panel represents the linear trendline (line of best fit) of the data, and dash lines represent 95% confidence interval.

To examine the effects of abstinence from oxycodone self-administration on neuronal structure in NAcSh, we analyzed 45 reconstructed MSNs from 23 female rats (8-14 cells from 5-7 rats among treatment groups) and 41 reconstructed MSNs from 21 male rats (6-16 cells from 3-8 rats among treatment groups). We found that soma sizes of the MSNs, estimated by maximal projection area at a single optical plane, vary in a fivefold range (∼40-200 µm2) in both male and female rats with identical cumulative frequency distributions (p = 0.954, Mann-Whitney test; **Figure 6D**). The varied soma sizes are correlated with dendritic size (dendritic length or volume), and, to a lesser degree, with the number of branch points (**Figure 6E, F**). This observation indicates a wide variation in the somatodendritic morphology of NAcSh-MSNs in both male and female rats. We then examined cumulative frequency distributions of soma size among four treatment groups, including male and female rats, and found no differences at 1 day of abstinence (p = 0.2703) or 14 days of abstinence (p = 0.5982, Kruskal-Wallis test, **Figure 6-Supplementary Figure 1B and F**). We did not detect significant effect of oxycodone abstinence on any parameter assayed in dendritic morphology (Two-way ANOVA, variables: treatment x sex) either at 1-day abstinence (F(1,40) = 0.455, p = 0.504 for dendritic length; F_(1,40)_ = 0.465, p = 0.499 for branch points), or at 14-days abstinence (F_(1,38_) = 0.001, p = 0.972 for dendritic length; F_(1,38)_ = 0.114, p = 0.738 for branch points, left panels in **Figure 6-Supplementary Figure 1C,D,G, H**). Consistent with these observations, Sholl analyses (right panels in **Figure 6-Supplementary Figure 1C,D,G,H**; three-way ANOVA, variables: treatment x sex x radial distance) indicate no changes after 1 day abstinence (F_(1, 40)_ = 0.4568, p = 0.5030 for dendritic length; F _(1, 40)_ = 1.547 p = 0.2208 for branch points), or at 14 days abstinence (F(_1, 38)_ = 0.001, p = 0.9693 for dendritic length; F_(1, 38)_ = 0.1138 p = 0.7378 for branch points). On the other hand, we observed small sex differences in the number of branch points (F_(1,40)_ = 4.444, p = 0.041; Bonferroni’s multiple comparisons test, p = 0.546 for male versus female saline rats) in NAcSh-MSNs at 1 day abstinence (whereas the effect on dendritic length was not significant, F_(1,40)_ = 3.396, p = 0.073; left panels in **Figure 6-Supplementary Figure 1C, D**). At 14 days of abstinence (left panels in **Figure 6-Supplementary Figure 1G,H**), there were no differences between male and female rats (F_(1,38)_ = 1.675, p = 0.203 for dendritic length; F_(1,38)_ = 1.964, p = 0.169 for branch points). No significant interaction between treatment and sex was found on dendritic length and branch points. The Sholl analyses of same data as above showed no sex differences both at 1 day abstinence (F_(1, 40)_ = 3.396, p = 0.0728 for dendritic length; F_(1, 40)_ = 1.848, p = 0.1816 for branch point) and at 14 days abstinence (F_(1, 38)_ = 1.670, p =0.2041 for dendritic length; F_(1, 38)_ = 1.964, p = 0.1692 for branch point). Taken together, our quantitative analyses demonstrate the lack of effect on somatodendritic morphology of NAcSh-MSNs at either 1 day or 14 days of abstinence from oxycodone self-administration in both female and male rats.

## Discussion

In this study, we investigated sex differences in withdrawal symptoms, cue-induced oxycodone-seeking (relapse), and changes in PVT-NAcSh synaptic transmission following short and long periods of abstinence from oxycodone self-administration. Our results indicate that acute abstinence increased withdrawal symptoms and cue-induced oxycodone-seeking without affecting PVT-NAcSh glutamatergic transmission, and no significant sex differences were observed. In contrast, prolonged abstinence increased cue-induced oxycodone-seeking and the efficacy of synaptic transmission in PVT-NAcSh projections in both male and female rats, with a stronger effect on cue-induced oxycodone-seeking in females.

### Oxycodone self-administration is similar between male and female rats

Escalation of drug taking in rats is defined as an increase in the amount and frequency of drug intake over time during self-administration protocols. It is used as a model of addiction, specifically to study tolerance to drugs taken and dependence on them. Our findings revealed that escalation of intravenous oxycodone self-administration under long access conditions was similar between female and male rats, indicating that sex does not significantly affect the development of oxycodone addiction using this protocol of self-administration. This is consistent with our previous observations when long-access self-administration was used [28,33]. However, it contrasts with other studies that have found that female rats orally self-administered more oxycodone than males in a short-access (1-hr) FR-1 schedule self-administration paradigm [39], whereas other studies from our lab have shown that males intravenously self-administered more oxycodone compared to females in the first three trials of an FR-1 schedule of reinforcement [25]. These results suggest that specific behavioral measurements and routes of administration may reveal differences in the development and maintenance of oxycodone use in male and female subjects.

### Increased spontaneous withdrawal symptoms after acute abstinence in males and females

Spontaneous withdrawal symptoms from opioids use refer to the symptoms that occur when an individual who has been taking opioids regularly suddenly stops using them or reduces their dose.

These symptoms occur as a result of the body’s physical dependence on the drug. Here, we found that both male and female rats exhibited increased spontaneous withdrawal signs after acute abstinence from oxycodone self-administration. These findings are consistent with the results of previous experiments with non-contingent morphine administration, which showed a similar increase in withdrawal symptoms after 1-day of abstinence in both males and females, with males showing a stronger effect [26]. However, we did not find a significant difference in the magnitude of spontaneous withdrawal signs between sexes following oxycodone self-administration in rats, highlighting the importance of considering the drug’s mechanism of action and administration routes when assessing its potential to affect males and females differently.

### Sex differences in cue-induced oxycodone-seeking after prolonged abstinence

In rats, abstinence periods from drug use have been shown to lead to the development of drug-seeking behaviors, which can intensify over time and lead to relapse. It has been shown that the longer abstinence periods result in a greater intensity of drug-seeking [10,40,41]. During this period, there are changes in the brain that can lead to the development of drug cravings, including alterations in synaptic function and changes in the expression of neurotransmitters and their receptors [10,23,42–46]

In addition, a recent study [47] found that male rats that underwent a longer abstinence period (30 days) following oxycodone self-administration showed significantly higher levels of drug-seeking behavior compared to rats that underwent a shorter abstinence period (1-day). This suggests that the incubation of oxycodone craving occurs in a time-dependent manner, with longer abstinence periods leading to increased drug-seeking behavior. Here we found that females but not males have an increased in cue-induced oxycodone-seeking after prolonged abstinence (14-days) compared to acute (1-day) abstinence. Females also exhibited increased drug-seeking during the prolonged abstinence period compared to males. This suggests that cue-induced cravings for oxycodone may develop faster and they are stronger in females compared to males.

### PVT-NAcSh synaptic strength increases after prolonged abstinence in both male and female rats

Furthermore, we investigated the effects of abstinence on glutamatergic synaptic transmission in PVT-NAcSh projections. Both the PVT and the NAc are highly heterogeneous nuclei with distinctive anatomical subregions (anterior/posterior PVT; core/shell NAc) that contain different types of cells (Type I and Type II in the PVT; dopamine receptor 1 (D1)- and dopamine receptor 2 (D2)-expressing medium spiny neurons (MSNs; D1-MSN or D2-MSN) in the NAc). Recent studies have highlighted the role of the PVT in modulating reward-seeking behaviors, particularly in response to drugs of abuse [48–52]. Several studies have shown that the PVT is involved in retrieving and consolidating drug-associated memories [53,54]. For example, one study found that optogenetic activation of PVT inputs to the NAc was sufficient to reinstate drug-seeking behavior in rats [15]. Another study showed that silencing PVT neurons during the retrieval of opiate-associated memories impaired the expression of drug-seeking behavior [53]. The PVT has also been implicated in the regulation of drug-seeking behavior more broadly. For example, transient inactivation of the posterior PVT can block cocaine-seeking behavior in rats [50], which seems to be dependent on the type of reinstatement model and individual differences [55]. Additionally, the PVT has been shown to play a key role in cue-induced drug-seeking after abstinence [13], suggesting that it serves an important function in the neural circuitry underlying relapse.

Here, we found that acute abstinence from oxycodone self-administration did not appear to affect neurotransmission in the PVT-NAcSh projections, as assessed by the EPSC amplitude input-output curves and paired-pulse ratio. Furthermore, the rectification index for the AMPAR-mediated EPSCs and AMPAR/NMDAR EPSC amplitude ratio at the PVT-NAcSh synapses were not significantly different between treatment groups or sexes, indicating that subunit composition of postsynaptic AMPA receptors or their sensitivity to glutamate were not affected by acute abstinence from oxycodone self-administration. These results suggest that a brief period of abstinence may not be sufficient to induce significant changes in synaptic properties of the PVT-NAcSh circuit.

In contrast, prolonged abstinence from oxycodone self-administration resulted in significant changes in glutamatergic synaptic transmission in the PVT-NAcSh circuits, as measured by EPSC input-output curves and paired-pulse ratio. Specifically, prolonged abstinence resulted in an increase in EPSC amplitude and a decrease in the paired-pulse ratio, suggesting that there is an increase in the probability of neurotransmitter release and/or alterations in presynaptic release mechanisms in the studied pathway. However, prolonged abstinence did not change the rectification index of the EPSCs or the AMPAR/NMDAR EPSC amplitude ratio, indicating that there is no change in the subunit composition of AMPA receptors at the PVT-NAcSh glutamatergic synapses.

These results suggest that long-term abstinence from oxycodone self-administration is associated with significant changes in synaptic properties of the PVT-NAcSh circuit—presynaptically-expressed synaptic potentiation, specifically—potentially contributing to the persistent changes in reward processing and relapse vulnerability that are often observed in individuals with opioid use disorder.

### Lack of changes in intrinsic excitability of MSNs in NAcSh after prolonged abstinence

The NAc plays an important role in regulating motivated behaviors such as reward-seeking and cravings [56,57]. Within the NAc, MSN intrinsic excitability regulates NAc responses to afferent inputs and controls the NAc outputs to other components of reward system in the brain. In our study, we investigated changes in NAcSh MSN intrinsic excitability following acute or prolonged abstinence from oxycodone self-administration using nested hierarchical Poisson GLMMs to appropriately account for the hierarchical structure of the electrophysiological data.

One goal of this study was to examine the interplay between PVT-NAcSh glutamatergic transmission and NAcSh MSN intrinsic excitability following oxycodone abstinence. The standing theory suggests that MSN excitability decreases as a homeostatic response to increased glutamatergic input [23,43,58]. Our data do not support a compensatory decrease in excitability in either sex at either abstinence time point. Instead, the trend toward increased excitability in males after prolonged abstinence, if confirmed with adequate sample sizes, would suggest that prolonged oxycodone abstinence may produce maladaptive neuroadaptations that enhance rather than suppress MSN excitability. Such an increase, acting in concert with the strengthening of PVT-NAcSh synaptic transmission observed in the current study, could facilitate signal flow through this circuit and enhance MSN output to downstream targets, potentially contributing to relapse-related behaviors.

Future studies with larger cohorts will be necessary to determine whether this trend in males represents a reproducible and sex-specific neuroadaptation following prolonged oxycodone abstinence.

### Implications of anatomical and functional distinctions in PVT-NAcSh circuits for relapse

Our study was designed to target posterior PVT-NAcSh projections, based on previous studies [13,16,21]. The anterior/posterior PVT have been shown to have different subcortical targets [59] and increasing evidence demonstrates substantial functional differences in anterior vs posterior PVT outputs [60]. Posterior projections to the NAc are selectively tuned to aversive stimuli, while the anterior PVT is more involved in reward-seeking behaviors [61]. Furthermore, our goal was to determine the overall effects of acute versus prolonged abstinence on synaptic and neuronal properties in PVT-NAcSh circuits in male and female rats, as opposed to differentiating between the contributions of D1 and D2 receptor-expressing MSNs. However, recent studies demonstrated differences in the roles of PVT-to-D1, PVT-to-D2 MSN, and PVT-to-Interneurons in mediating drug-seeking or withdrawal after either acute or prolonged abstinence from opioid self-administration [13,21]. Thus, it would be interesting to explore, in future studies, the effects of our oxycodone administration regimens on neurotransmission in PVT projections to D1- or D2-MSNs, using *ex vivo* optogenetic studies analogous to those described in the present work.

Previous work has demonstrated cell-type-specific plasticity at PVT-NAc circuit following heroin self-administration and extinction. It includes strengthening PVT inputs to D2 MSNs and reducing synaptic strength from PVT to parvalbumin-expressing interneurons (PV-INs) [21]. In more detail, stimulation of PVT terminals in the NAcSh has been shown to increase opioid-seeking prior to, but not after, extinction of drug availability[13]. Consistent with this behavioral shift, extinction training can occlude the increase in synaptic strength at PVT to NAc D2 MSNs observed following heroin self-administration in mice [21]. This increase in synaptic strength was reflected by enhanced opto-evoked EPSC amplitude, an increased AMPA/NMDA ratio, and a decreased paired-pulse ratio (PPR), all consistent with potentiated excitatory transmission. PVT projections also synapse onto parvalbumin-expressing interneurons in the NAc (NAc PV-INs; decreased opto-evoked EPSCs and AMPA/NMDA ratios) [21], where an overall decrease in synaptic strength has been reported after both self-administration and extinction [21]. In contrast, PVT-to-NAc D1 MSNs do not show changes in synaptic strength during self-administration or extinction [21]. However, a shift in rectification index following heroin self-administration suggests increased calcium-permeable AMPA receptor (CP-AMPAR) function at these synapses [21]. Importantly, restoring normal synaptic transmission in the PVT-to-NAc PV-IN pathway was sufficient to prevent relapse-like behavior, measured as active lever pressing for heroin [21]. Furthermore, a recent study found that prolonged abstinence period increased CP-AMPAR expression in D1 and D2 MSNs in the NAc of male and female rats [62].

In a morphine conditioned place preference (CPP) model, transient inhibition of the PVT-NAc pathway prevented morphine-primed relapse after both short (4 days) and prolonged (14 days) abstinence without affecting morphine-induced locomotor activity. Importantly, inhibition of this pathway disrupted relapse only when animals were re-exposed to the drug-associated context, indicating that PVT-NAc activity is required for the retrieval and maintenance of opiate-associated memories [53].

Consistent with a circuit architecture in which PVT inputs both directly excite MSNs and recruit local inhibitory microcircuits, our results show that stimulation of glutamatergic PVT terminals in the NAcSh evokes monosynaptic excitation alongside disynaptic inhibition mediated by local GABAergic interneurons. Following prolonged abstinence from oxycodone, excitatory synaptic transmission at PVT-NAcSh projections was strengthened, suggesting enhanced excitatory drive within this pathway. In contrast, inhibitory synaptic transmission was not significantly altered in either males or females, indicating that feedforward inhibitory recruitment and overall inhibitory tone within the PVT-NAcSh circuit remain intact following prolonged abstinence from oxycodone self-administration. Together, these findings suggest that prolonged oxycodone abstinence selectively potentiates excitatory rather than inhibitory synaptic transmission within this pathway, potentially shifting the excitation-inhibition balance in a manner that may contribute to relapse-related behaviors.

In our study, morphological analyses revealed no detectable changes in dendritic length or branching complexity in NAcSh MSNs at either 1 day or 14 days of abstinence following oxycodone self-administration. Additionally, no sex differences were observed in the somatodendritic morphology of NAcSh MSNs between control male and female rats. These findings suggest that the abstinence-dependent adaptations previously reported in this circuit may occur primarily through synaptic or molecular mechanisms, rather than through large-scale structural remodeling of MSN dendritic architecture.

## Conclusion

When considered alongside prior work, these results support a model in which the anatomical and cell-type-specific organization of PVT inputs to the NAcSh plays a critical role in relapse-related behaviors. Specifically, PVT projections onto D2 MSNs and PV interneurons may contribute to relapse vulnerability during early or acute abstinence, potentially through mechanisms linked to negative affective states. In contrast, adaptations involving D1 MSNs, including CP-AMPAR recruitment during prolonged abstinence, may underlie the incubation of craving that emerges after extended drug abstinence.

Overall, our findings provide further insight into the circuit mechanisms underlying oxycodone relapse and highlight the importance of considering cell-type specificity, abstinence duration, and behavioral context when interpreting plasticity within the PVT-NAc pathway. Importantly, differences across studies may also reflect variations in drug exposure paradigms, including contingent (self-administration) versus non-contingent drug delivery, as well as whether animals undergo extinction training or remain in forced abstinence. These experimental variables engage distinct learning processes and motivational states that can differentially shape circuit adaptations and relapse-related behaviors.

Future studies should aim to dissect the distinct contributions of anterior versus posterior PVT projections onto D1 and D2 MSNs in the NAcSh and determine how selective manipulation of these pathways influences relapse behaviors across different abstinence states and relapse triggers (e.g., drug-, cue-, or stress-induced). In addition, understanding how thalamic inputs interact with cortical projections to ventral striatal circuits during abstinence and relapse represents an important avenue for future investigation.

## Methods

### Subjects

Adult male (250-275g; Total N = 42) and female (200-225g; Total N = 45) Sprague Dawley rats (Charles River Laboratory, Wilmington, MA) were used in this study. Upon arrival, rats were group housed (4 rats per cage) and were habituated for 1 week to the animal colony kept on a 12-h light/dark cycle (lights on 7:00AM) with food and water *ad libitum*. Following surgeries, rats were singly-housed for the rest of the experiment. All animal procedures were conducted in accordance with the guidelines of the National Institutes of Health and were approved by the Institutional Animal Care and Use Committee (IACUC) at McLean Hospital/Harvard Medical School (Protocol #2017N0000277, #2018N000090 & #2017N000125).

### Surgeries

#### Stereotaxic surgery and viral injections

All surgeries were performed according to AAALAC guidelines. Rats were first anesthetized with ketamine and xylazine (80 mg/kg and 8 mg/kg, respectively,I.P.). A craniotomy was made to target the PVT using the following stereotaxic coordinates, based on Paxinos and Watson 6th edition rat brain atlas [24]: AP: -2.6mm from Bregma, ML: +2.0mm from Bregma (20° angle ML), and DV: -6.3mm from skull surface at injection site. A total of 1μL of the AAV5 vector carrying CaMKIIα-ChR2(H143R)-eYFP was injected unilaterally into PVT at a rate of 125 nl/min using a 10μl Hamilton syringe with a 29-gauge needle under the control of a micro-syringe pump (Harvard Instruments). Viral vectors (titers, ∼10.0 x 10^12^ particles/ml) were purchased from the University of North Carolina viral vector facility.

#### Intravenous catheter implantation surgery

After recovery from stereotaxic surgery (∼7 days), rats were implanted with indwelling silastic intravenous jugular catheters (SAI infusions; RSB-SA-7.5CF and RSB-SA-7.5CM), as described in [25–27]. Rats were anesthetized with ketamine and xylazine (80 mg/kg and 8 mg/kg, respectively, I.P.), and catheters were implanted into the right jugular vein, secured to the vein with non-absorbable suture thread and passed subcutaneously through the rat’s back. All rats received an injection of ketofen (5mg/kg; S.C.) and gentamicin (0.1ml; 10mg/ml, I.V.) during catheter implantation. Catheters were flushed daily with 0.2ml of heparinized saline (30 units/ml; I.V.) and once a week with 0.2ml of gentamicin (10 mg/ml I.V.). Catheter patency was checked once per week using methoexital (Brevital; 0.1 ml females; 0.2 males of 10 mg/ml I.V.).

### Behavioral methods

#### Oxycodone self-administration

Med Associates operant conditioning chambers (30.5 (*l*) × 24.1 (*w*) × 29.2 (*h*) cm), kept within soundproofed outer chambers with ventilation fans, were equipped with two retractable levers, each with a cue light above them, a house light, a counterbalance swivel and tether, and an infusion pump. Rats (males: n = 15 saline, 27 oxycodone; females: n= 22, saline, 23 oxycodone) underwent 8 days of short access (ShA) oxycodone self-administration training (0.06mg/kg/infusion; 1h/day) followed by 14 days of long access (LgA) regimen (0.06 mg/kg/infusion; 6h/day), similar to Mavrikaki et al 2019 [28]). Self-administration sessions were run 7 days/week, at approximately 9:00 am each day. All self-administration was conducted during the light phase of a 12:12 light/dark cycle (lights on at 7:00am; lights off at 7:00pm). A fixed-ratio 1 (FR1) schedule of reinforcement was used such that a press on the active lever resulted in a 4-s oxycodone infusion (100 µl) followed by a 6-s time out period where a press on the active lever produced no consequences.

#### Assessing oxycodone dependence

To demonstrate that our oxycodone self-administration protocol induced dependence in both male and female rats, we measured spontaneous somatic withdrawal signs 24-h after the last oxycodone self-administration session. After removal from self-administration chambers and catheter flushing, rats were placed back in their home cages and brought to a quiet, temperature-maintained (20°C) room and allowed to habituate for ∼15-min. Rats were then individually placed into clear, 65-cm-high by 25-cm-diameter Plexiglas cylinders that contained a small amount of bedding. Rats were allowed to habituate to the cylinders for ∼15 min. At this point, a digital video system (Swann Communications, Sante Fe, CA) was used to record the rats in the cylinders for 20 minutes. Upon completion of recording, somatic withdrawal behaviors were scored for the first 15 min of the recording by a researcher who was unaware of the treatments. Every 15 seconds, the following behaviors were marked as either present or absent: diarrhea, ptosis, jumping, walking, rearing, digging, flat posture, “wet dog shakes,” grooming and teeth chattering [26,29]. The number of occurrences of each behavior was summed. In addition, a Total Withdrawal Score was calculated by summing weighted frequencies of those behaviors most commonly and specifically observed in opioid withdrawal: Total Withdrawal = Grooming (x1.0) + Wet Dog Shakes (x1.5) + Ptosis (x1.2) [29,30]. Wet Dog Shakes and Ptosis were multiplied by previously determined weighting factors to account for their high importance, but low prevalence, to withdrawal signs.

#### Forced abstinence and cue-induced oxycodone-seeking

a. *1-day abstinence* (acute abstinence): From the total rats above, male (Saline, N = 9; Oxycodone, N = 18) and female (Saline, N = 13; Oxycodone, N = 15) rats underwent 1d of forced abstinence (rats returned to the vivarium in their home cages for 24-h) from oxycodone self-administration. A subset of 1d abstinence male (Saline, N=5; Oxycodone N=10) and female (Saline, N=7; Oxycodone N=7) rats were used to measure somatic withdrawal signs, and another subset of males (Saline, N=4; Oxycodone, N=8) and females (Saline, N=6; Oxycodone, N=8) was used to measure cue-induced oxycodone-seeking after the 1d abstinence period. After 1d of abstinence, rats were reintroduced to the operant chamber for a 2-h relapse test. The cues associated with oxycodone were presented, but no drug was delivered.
b. *14-day abstinence* (prolonged abstinence): From the total rats above, male (Saline, N=6; Oxycodone N=8) and female (Saline, N=8; Oxycodone, N=7) rats underwent 14d of abstinence from oxycodone self-administration. After 14d of abstinence, rats were reintroduced to the operant chambers for a 2-h relapse test as described above.

The number of active and inactive lever presses were recorded during incubation/ cue-induced oxycodone-seeking testing following acute and prolonged abstinence periods. Active lever presses were compared between saline and oxycodone groups for cue-induced oxycodone-seeking. Rat brains were extracted thirty minutes to one hour following the cue-induced oxycodone-seeking test, such that electrophysiological recordings reflect synaptic properties associated with relapse after acute or prolonged abstinence.

### Ex-vivo electrophysiology and optogenetic stimulation

Coronal slices (300 µm in thickness) containing the NAc were obtained using a vibratome in cold cutting solution containing the following in mM: 252.0 sucrose, 1.0 CaCl_2_, 5.0 MgCl_2_, 2.5 KCl, 1.25 NaH_2_PO_4_, 26.0 NaHCO_3_ and 10.0 glucose and equilibrated with 95% O_2_ and 5% CO_2_. Slices were then incubated in artificial cerebrospinal fluid (ACSF) containing the following in mM: 125 NaCl, 2.5 KCl, 2.5 CaCl_2_, 1.0 MgSO_4_, 1.25 NaH_2_PO_4_, 26.0 NaHCO_3_, and 10.0 glucose at room temperature for at least 1 hr before recordings started. Whole-cell recordings were obtained from the NAcSh neurons with patch electrodes (3-5 MΩ resistance) containing the following in mM: 135.0 Cs-methane-sulfonate, 5.0 NaCl, 1.0 MgCl_2_, 10.0 BAPTA, 10.0 HEPES, 2.0 ATP and 0.20 GTP adjusted to pH 7.2 with CsOH. Neurobiotin (0.2%; Vector Laboratories) was also added to the internal solution before the recordings to allow subsequent histochemical localization of the recorded neurons in the NAcSh.

Synaptic responses were induced by photostimulation of ChR2-expressing PVT projecting terminals in the NAcSh with a LED light source (excitation wavelength: 470 nm, 5 ms in duration, Thorlabs). Whole-cell recordings were accepted if the access resistance was ≤20 MΩ and remained stable throughout the recording period (defined as <20% change from baseline). Recordings that exceeded these thresholds were excluded from analysis. All recordings were performed at 30-32°C. After recordings, slices were placed in PBS containing 4% paraformaldehyde and kept in the refrigerator until histological processing.

The pharmacological reagents used in electrophysiological experiments included NBQX disodium salt, D-AP5, NBQX, and (-)-bicuculline methobromide, which were prepared as stock solutions in water at 1000- to 5000-fold concentrations and stored at -20°C.

### Morphological Analysis

#### Histology for Neurobiotin-filled cells

Brain slices containing Neurobiotin-filled neurons in NAcSh were washed in PBS for 20 min x 3 times and incubated with Streptavidin Alexa 568 conjugate (10-20 µg/ml, catalog number: S11226, Molecular Probes) in PBS containing 0.2% Triton X-100 at room temperature for 24 hours. The slices were then washed with PBS for 20 min x 3 times and mounted on gelatinized slides. The anti-fading mounting media with DAPI (Vectashield, Vector Laboratories) was applied to slices.

#### Analysis of dendritic morphology of NAcSh-MSNs

Acquisition of imaging data of Neurobiotin-stained neurons was performed using a Leica SP8 TCS confocal microscope under a 40X/1.30 NA oil-immersion objective lens. The image resolution (1024 X 1024 pixels) and z step (0.5 μm) of optical planes were kept constant throughout the study. To image entire dendritic trees of NAcSh neurons, the zoom was adjusted in a range of 0.75 – 1.0, which corresponds to the voxel size between 0.284 X 0.284 X 0.5 μm and 0.379 X 0.379 X 0.5 μm. Three-dimensional (3D) reconstruction of dendritic trees was conducted in stacked confocal images in the program NeuronStudio (version 0.9.92, Icahn School of Medicine at Mount Sinai, New York, NY; [31].

We used the manual tracing tool to reconstruct dendrites starting from the soma. Specifically, we started the manual tracing from the beginning of each primary dendrite and moved one node at a time to form an entire path along a branch through the views at XY, ZY and XZ orientations. The program provides numerical measurements of the reconstructed dendritic trees, including dendritic length, surface area, volume, number of branch points, and Sholl analysis. Soma size was estimated by measuring the maximal projection area of a neuron soma in a single optical coronal section (1.038 µm thick) using ImageJ. The spatial organization of the dendritic field was characterized by the ratio of the long axis to the short axis of the dendritic field at the coronal plane. The long axis of the dendritic field was defined as the distance from the soma to the most distal dendritic process, and the short axis as the distance from the soma to the most distal dendritic point located 90° from the long axis [32]. The measurements were performed on stacked images of MSNs using the Leica Application Suite X (LAS X, Leica Microsystems CMS GmbH, version 3.5.5). The Neurobiotin-stained neurons were primarily obtained from the medial division of NAcSh in two adjacent slices of caudal nucleus accumbens, corresponding to the bregma levels between 10.08 and 10.56 mm [24].

### Statistical Analysis

Male and female rats were randomly assigned to either saline or oxycodone groups. In electrophysiological experiments, ∼2-3 neurons were recorded per animal. The numbers of rats and recorded neurons for the analysis of the different experiments are indicated in the results section. Data are reported as mean +/- SEM. All electrophysiology data were collected using Patch Master (Heka systems). We used Prism 9 (GraphPad) for statistical analysis using Two-tailed t-tests, Mixed effects model (Restricted Maximum Likelihood/REML), and Two- or Three-way ANOVAs with Sidak’s multiple comparisons, as appropriate.

For analyses of intrinsic excitability based on spike count–current relationships, spike output (number of action potentials per current step) was analyzed using nested hierarchical Poisson generalized linear mixed-effects models (GLMMs), fitted separately for each sex and abstinence duration. Each model included injected current (mean-centered), drug condition (Saline vs. Oxycodone), and their interaction as fixed effects. Random effects included random intercepts and slopes for injected current at the animal level, and random intercepts for cells nested within animals, to account for the non-independence of repeated current steps within cells and of multiple cells recorded from the same animal. Unique identifiers were assigned to each animal and cell by concatenating condition labels with animal and cell IDs, ensuring that animals across treatment conditions were treated as independent subjects. Models were fitted using maximum likelihood estimation with Laplace approximation. Overdispersion was assessed using the Pearson chi-squared statistic divided by the residual degrees of freedom; when this ratio exceeded 1.5, a dispersion correction was applied within the Poisson framework. Statistical significance of fixed effects was assessed using marginal F-tests with residual degrees of freedom. Sex comparisons among oxycodone-treated animals were conducted as separate analyses using identical model structures, with sex as the grouping condition. For visualization, marginal model-predicted values were overlaid on each plot as dashed lines, representing the population-level fixed-effects predictions from the fitted GLMM with random effects set to zero, back-transformed from the log scale to spike counts. To assess statistical power, a posthoc power analysis was conducted for each comparison using a two-sample t-test approximation at the rat level, with rat-level mean spike output as the unit of analysis and the pooled between-rat standard deviation as the denominator for effect size estimation (Cohen’s d). This approach was used to estimate the number of animals per group required to achieve 80% and 90% statistical power, given the observed effect sizes. All analyses were performed in MATLAB (R2025b).

IPSC amplitudes were analyzed using linear mixed-effects models (LMEs) to account for the hierarchical structure of the data, with repeated measurements across light intensities nested within animals and multiple cells recorded per animal. Unique identifiers were assigned to each rat and cell by concatenating condition labels with animal and cell IDs, ensuring that animals across treatment conditions were treated as independent subjects. Models included fixed effects of light intensity (mean-centered across the six stimulation levels: 0.5, 1.3, 2.0, 4.7, 7.6, and 10.4 mW), treatment condition (Saline vs. Oxycodone), and their interaction. Random effects included random intercepts and light intensity slopes for individual animals, and random intercepts for cells nested within animals, capturing both between-animal variability in baseline responses and sensitivity to light, as well as within-animal variability across cells. Model parameters were estimated using restricted maximum likelihood (REML). Statistical significance of fixed effects was assessed using marginal F-tests with residual degrees of freedom, applied consistently across male and female datasets. Data are presented as mean ± SEM, computed from rat-level averages, in which responses from multiple cells within the same animal were first averaged to yield one value per animal per light intensity, consistent with the animal-level inference of the LME. Male and female datasets were analyzed separately using identical model structures to assess sex-specific effects. All analyses were performed in MATLAB (R2025b).

The number of animals and cells used per experiment, as well as the results of all statistical analyses, are reported in the text and figure legends.

## Acknowledgements

The authors would like to extend their gratitude to Dr. Cristina Berciu. This manuscript is dedicated to the memory of Dr. Elena H. Chartoff, who passed away on May 23, 2024. We honor Elena’s significant contributions to the field of behavioral sex differences and her unwavering support for young scientists and underrepresented minorities.

---

**Figure 6-Supplementary Figure 1.**
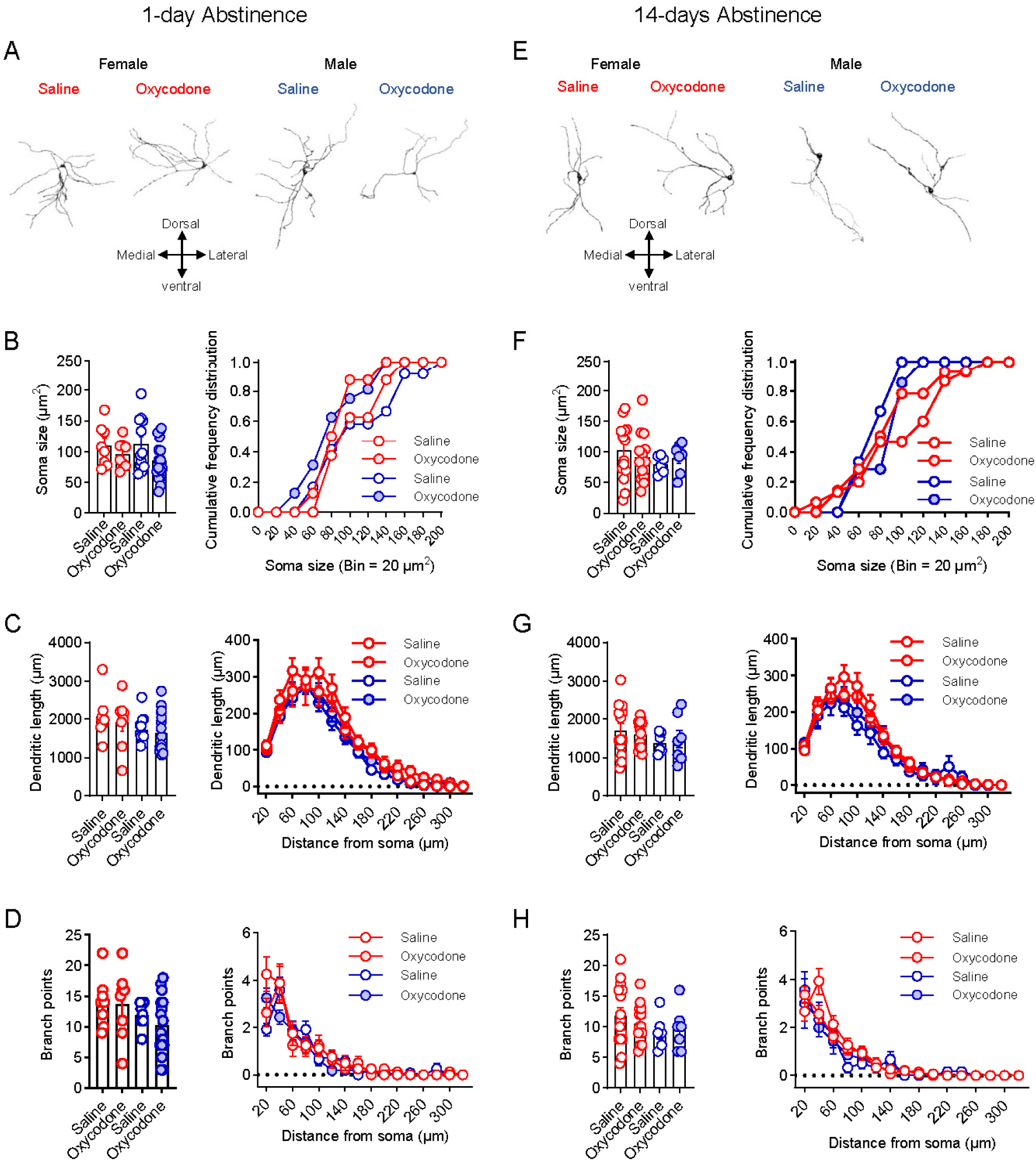
Dendritic morphology of NAcSh-MSNs remained unchanged at either 1-day or 14-days abstinence from oxycodone self-administration in both female and male rats. **A & E**: Representative reconstructed NAcSh-MSNs from saline or oxycodone group in female (**left**, 8-14 cells from 5-7 rats among four treatment groups) and male (**right**, 6-16 cells from 3-8 rats among four treatment groups) rats at 1-day (**A**) or 14-days (**E**) abstinence, respectively. **B & F**: There is no difference in soma sizes (**left**, p = 0.077 or 0.352, respectively, two-way ANOVA) and their cumulative frequency distributions (**right**, p = 0.270 or 0.598, respectively, Kruskal-Wallis test) among treatment groups at 1 day (**B**) or 14 days (**F**) abstinence after oxycodone self-administration. **C & G**: No difference was observed in the total dendritic length (**left**, p = 0.504 or 0.972, respectively, two-way ANOVA) and Sholl analysis on dendritic length (**right**, p = 0.503 or 0.969, respectively, three-way ANOVA) between saline and oxycodone groups at 1-day or 14-days abstinence. **D & H**: No difference was observed in the total number of branch points (**left**, p = 0.499 or 0.738, respectively) and Sholl analysis on branch points (**right**, p = 0.221 or 0.738, respectively) between saline and oxycodone group at 1-day or 14-days abstinence. **Red and blue symbols** represent data sets from **female or male** rats, respectively. **Open or filled** symbols represent the data set of **saline or oxycodone**, respectively.

## References

[1] Feltenstein MW, See RE, Fuchs RA. Neural Substrates and Circuits of Drug Addiction. Cold Spring Harb Perspect Med 2021;11:a039628. 10.1101/cshperspect.a039628.

[2] Strang J, Volkow ND, Degenhardt L, Hickman M, Johnson K, Koob GF, Marshall BDL, Tyndall M, Walsh SL. Opioid use disorder. Nat Rev Dis Primer 2020;6:1–28. 10.1038/s41572-019-0137-5.

[3] Golden SA, Jin M, Shaham Y. Animal Models of (or for) Aggression Reward, Addiction, and Relapse: Behavior and Circuits. J Neurosci Off J Soc Neurosci 2019;39:3996–4008. 10.1523/JNEUROSCI.0151-19.2019.

[4] Koob GF, Volkow ND. Neurocircuitry of addiction. Neuropsychopharmacol Off Publ Am Coll Neuropsychopharmacol 2010;35:217–38. 10.1038/npp.2009.110.

[5] Self DW. Neural substrates of drug craving and relapse in drug addiction. Ann Med 1998;30:379–89. 10.3109/07853899809029938.

[6] Self DW, Nestler EJ. Relapse to drug-seeking: neural and molecular mechanisms. Drug Alcohol Depend 1998;51:49–60. 10.1016/s0376-8716(98)00065-9.

[7] Venniro M, Caprioli D, Shaham Y. Animal models of drug relapse and craving: From drug priming-induced reinstatement to incubation of craving after voluntary abstinence. Prog. Brain Res., vol. 224, 2016, p. 25–52. 10.1016/bs.pbr.2015.08.004.

[8] Grimm JW, Hope BT, Wise RA, Shaham Y. Incubation of cocaine craving after withdrawal. Nature 2001;412:141–2. 10.1038/35084134.

[9] Lu L, Grimm JW, Dempsey J, Shaham Y. Cocaine seeking over extended withdrawal periods in rats: Different time courses of responding induced by cocaine cues versus cocaine priming over the first 6 months. Psychopharmacology (Berl) 2004;176:101–8. 10.1007/s00213-004-1860-4.

[10] Pickens CL, Airavaara M, Theberge F, Fanous S, Hope BT, Shaham Y. Neurobiology of the incubation of drug craving. Trends Neurosci 2011;34:411–20. 10.1016/j.tins.2011.06.001.

[11] Reiner DJ, Fredriksson I, Lofaro OM, Bossert JM, Shaham Y. Relapse to opioid seeking in rat models: behavior, pharmacology and circuits. Neuropsychopharmacology 2019;44:465–77. 10.1038/s41386-018-0234-2.

[12] Venniro M, Reverte I, Ramsey LA, Papastrat KM, D’Ottavio G, Milella MS, Li X, Grimm JW, Caprioli D. Factors modulating the incubation of drug and non-drug craving and their clinical implications. Neurosci Biobehav Rev 2021;131:847–64. 10.1016/j.neubiorev.2021.09.050.

[13] Giannotti G, Gong S, Fayette N, Heinsbroek JA, Orfila JE, Herson PS, Ford CP, Peters J. Extinction blunts paraventricular thalamic contributions to heroin relapse. Cell Rep 2021;36:109605. 10.1016/j.celrep.2021.109605.

[14] Peters J, Kalivas PW, Quirk GJ. Extinction circuits for fear and addiction overlap in prefrontal cortex. Learn Mem 2009;16:279–88. 10.1101/lm.1041309.

[15] Zhu Y, Wienecke CFR, Nachtrab G, Chen X. A thalamic input to the nucleus accumbens mediates opiate dependence. Nature 2016;530:219–22. 10.1038/nature16954.

[16] Zhu Y, Nachtrab G, Keyes PC, Allen WE, Luo L, Chen X. Dynamic salience processing in paraventricular thalamus gates associative learning. Science 2018;362:423–9. 10.1126/science.aat0481.

[17] Floresco SB, McLaughlin RJ, Haluk DM. Opposing roles for the nucleus accumbens core and shell in cue-induced reinstatement of food-seeking behavior. Neuroscience 2008;154:877–84. 10.1016/j.neuroscience.2008.04.004.

[18] Floresco SB. The nucleus accumbens: An interface between cognition, emotion, and action. Annu Rev Psychol 2015;66:25–32. 10.1146/ANNUREV-PSYCH-010213-115159.

[19] Floresco SB, Montes DR, Tse MMT, Holstein M van. Differential Contributions of Nucleus Accumbens Subregions to Cue-Guided Risk/Reward Decision Making and Implementation of Conditional Rules. J Neurosci 2018;38:1901–14. 10.1523/JNEUROSCI.3191-17.2018.

[20] Klawonn AM, Malenka RC. Nucleus Accumbens Modulation in Reward and Aversion. Cold Spring Harb Symp Quant Biol 2018;83:119–29. 10.1101/sqb.2018.83.037457.

[21] Paniccia JE, Vollmer KM, Green LM, Grant RI, Winston KT, Buchmaier S, Westphal AM, Clarke RE, Doncheck EM, Bordieanu B, Manusky LM, Martino MR, Ward AL, Rinker JA, McGinty JF, Scofield MD, Otis JM. Restoration of a paraventricular thalamo-accumbal behavioral suppression circuit prevents reinstatement of heroin seeking. Neuron 2024;112:772–785.e9. 10.1016/j.neuron.2023.11.024.

[22] Carlezon WA, Thomas MJ. Biological substrates of reward and aversion: A nucleus accumbens activity hypothesis. Neuropharmacology 2009;56:122–32. 10.1016/j.neuropharm.2008.06.075.

[23] Hearing M, Graziane N, Dong Y, Thomas MJ. Opioid and Psychostimulant Plasticity: Targeting Overlap in Nucleus Accumbens Glutamate Signaling. Trends Pharmacol Sci 2018;39:276–94. 10.1016/j.tips.2017.12.004.

[24] Paxinos G, Watson C. The Rat Brain in Stereotaxic Coordinates: Hard Cover Edition. Elsevier; 2006.

[25] Mavrikaki M, Pravetoni M, Page S, Potter D, Chartoff E. Oxycodone self-administration in male and female rats. Psychopharmacology (Berl) 2017;234:977–87. 10.1007/s00213-017-4536-6.

[26] Mavrikaki M, Lintz T, Constantino N, Page S, Chartoff E. Chronic opioid exposure differentially modulates oxycodone self-administration in male and female rats. Addict Biol 2021;26:e12973. 10.1111/adb.12973.

[27] Thomsen M, Caine SB. Chronic Intravenous Drug Self-Administration in Rats and Mice. Curr Protoc Neurosci 2005;32. 10.1002/0471142301.ns0920s32.

[28] Mavrikaki M, Anastasiadou E, Ozdemir RA, Potter D, Helmholz C, Slack FJ, Chartoff EH. Overexpression of miR-9 in the Nucleus Accumbens Increases Oxycodone Self-Administration. Int J Neuropsychopharmacol 2019;22:383–93. 10.1093/ijnp/pyz015.

[29] Chartoff EH, Mague SD, Barhight MF, Smith AM, Carlezon WA. Behavioral and Molecular Effects of Dopamine D1 Receptor Stimulation during Naloxone-Precipitated Morphine Withdrawal. J Neurosci 2006;26:6450–7. 10.1523/JNEUROSCI.0491-06.2006.

[30] Chartoff EH, Barhight MF, Mague SD, Sawyer AM, Carlezon WA. Anatomically dissociable effects of dopamine D1 receptor agonists on reward and relief of withdrawal in morphine-dependent rats. Psychopharmacology (Berl) 2009;204:227–39. 10.1007/s00213-008-1454-7.

[31] Rodriguez A, Ehlenberger D, Kelliher K, Einstein M, Henderson SC, Morrison JH, Hof PR, Wearne SL. Automated reconstruction of three-dimensional neuronal morphology from laser scanning microscopy images. Methods 2003;30:94–105. 10.1016/s1046-2023(03)00011-2.

[32] O’Donnell P, Grace AA. Physiological and morphological properties of accumbens core and shell neurons recorded in vitro. Synapse 1993;13:135–60. 10.1002/syn.890130206.

[33] Guha S, Alonso-Caraballo Y, Driscoll GS, Babb JA, Neal M, Constantino NJ, Lintz T, Kinard E, Chartoff E. Ranking the contribution of behavioral measures comprising oxycodone self-administration to reinstatement of drug-seeking in male and female rats. Front Behav Neurosci 2022:470.

[34] Kalamarides DJ, Singh A, Wolfman SL, Dani JA. Sex differences in VTA GABA transmission and plasticity during opioid withdrawal. Sci Rep 2023;13:8460. 10.1038/s41598-023-35673-9.

[35] Cho JH, Deisseroth K, Bolshakov VY. Synaptic encoding of fear extinction in mPFC-amygdala circuits. Neuron 2013;80:1491–507. 10.1016/j.neuron.2013.09.025.

[36] Clem RL, Huganir RL. Calcium-Permeable AMPA Receptor Dynamics Mediate Fear Memory Erasure. Science 2010;330:1108–12. 10.1126/science.1195298.

[37] van Dongen YC, Mailly P, Thierry A-M, Groenewegen HJ, Deniau J-M. Three-dimensional organization of dendrites and local axon collaterals of shell and core medium-sized spiny projection neurons of the rat nucleus accumbens. Brain Struct Funct 2008;213:129–47. 10.1007/s00429-008-0173-5.

[38] Shi WX, Rayport S. GABA synapses formed in vitro by local axon collaterals of nucleus accumbens neurons. J Neurosci Off J Soc Neurosci 1994;14:4548–60. 10.1523/JNEUROSCI.14-07-04548.1994.

[39] Fulenwider HD, Nennig SE, Hafeez H, Price ME, Baruffaldi F, Pravetoni M, Cheng K, Rice KC, Manvich DF, Schank JR. Sex differences in oral oxycodone self-administration and stress-primed reinstatement in rats. Addict Biol 2020;25:e12822. 10.1111/adb.12822.

[40] Fuchs RA, Lasseter HC, Ramirez DR, Xie X. Relapse to drug seeking following prolonged abstinence: the role of environmental stimuli. Drug Discov Today Dis Models 2008;5:251–8. 10.1016/j.ddmod.2009.03.001.

[41] Zhou W, Zhang F, Liu H, Tang S, Lai M, Zhu H, Kalivas PW. Effects of training and withdrawal periods on heroin seeking induced by conditioned cue in an animal of model of relapse. Psychopharmacology (Berl) 2009;203:677–84. 10.1007/s00213-008-1414-2.

[42] De Vries TJ, Schoffelmeer ANM, Binnekade R, Raasø H, Vanderschuren LJMJ. Relapse to cocaine- and heroin-seeking behavior mediated by dopamine D2 receptors is time-dependent and associated with behavioral sensitization. Neuropsychopharmacology 2002;26:18–26. 10.1016/S0893-133X(01)00293-7.

[43] Dong Y, Taylor JR, Wolf ME, Shaham Y. Circuit and synaptic plasticity mechanisms of drug relapse. J Neurosci 2017;37:10867–76. 10.1523/JNEUROSCI.1821-17.2017.

[44] Loweth JA, Tseng KY, Wolf ME. Adaptations in AMPA receptor transmission in the nucleus accumbens contributing to incubation of cocaine craving. Neuropharmacology 2014;76:287–300. 10.1016/j.neuropharm.2013.04.061.

[45] McCutcheon JE, Loweth JA, Ford KA, Marinelli M, Wolf ME, Tseng KY. Group I mGlur activation reverses cocaine-induced accumulation of calcium-permeable AMPA receptors in nucleus accumbens synapses via a protein kinase C-dependent mechanism. J Neurosci 2011;31:14536–41. 10.1523/JNEUROSCI.3625-11.2011.

[46] Scheyer AF, Loweth JA, Christian DT, Uejima J, Rabei R, Le T, Dolubizno H, Stefanik MT, Murray CH, Sakas C, Wolf ME. AMPA Receptor Plasticity in Accumbens Core Contributes to Incubation of Methamphetamine Craving. Biol Psychiatry 2016;80:661–70. 10.1016/j.biopsych.2016.04.003.

[47] Wong B, Zimbelman AR, Milovanovic M, Wolf ME, Stefanik MT. GluA2-lacking AMPA receptors in the nucleus accumbens core and shell contribute to the incubation of oxycodone craving in male rats. Addict Biol 2022;27:e13237. 10.1111/adb.13237.

[48] Aurélie De Groote, Alban de Kerchove d’Exaerde. Thalamo-Nucleus Accumbens Projections in Motivated Behaviors and Addiction. Front Syst Neurosci 2021;15:711350. 10.3389/fnsys.2021.711350.

[49] Kirouac GJ. Placing the paraventricular nucleus of the thalamus within the brain circuits that control behavior. Neurosci Biobehav Rev 2015;56:315–29. 10.1016/j.neubiorev.2015.08.005.

[50] Matzeu A, Zamora-Martinez ER, Martin-Fardon R. The paraventricular nucleus of the thalamus is recruited by both natural rewards and drugs of abuse: recent evidence of a pivotal role for orexin/hypocretin signaling in this thalamic nucleus in drug-seeking behavior. Front Behav Neurosci 2014;8:117. 10.3389/fnbeh.2014.00117.

[51] Millan EZ, Ong ZY, McNally GP. Paraventricular thalamus: Gateway to feeding, appetitive motivation, and drug addiction. Prog. Brain Res., vol. 235, Elsevier B.V.; 2017, p. 113–37. 10.1016/bs.pbr.2017.07.006.

[52] Zhou K, Zhu Y. The paraventricular thalamic nucleus: A key hub of neural circuits underlying drug addiction. Pharmacol Res 2019;142:70–6. 10.1016/j.phrs.2019.02.014.

[53] Keyes PC, Adams EL, Chen Z, Bi L, Nachtrab G, Wang VJ, Tessier-Lavigne M, Zhu Y, Chen X. Orchestrating Opiate-Associated Memories in Thalamic Circuits. Neuron 2020;107:1113–1123.e4. 10.1016/j.neuron.2020.06.028.

[54] Pribiag H, Lim BK. Thalamic Retrieval of Opioid Memories. Neuron 2020;107:992–4. 10.1016/j.neuron.2020.09.006.

[55] Kuhn BN, Klumpner MS, Covelo IR, Campus P, Flagel SB. Transient inactivation of the paraventricular nucleus of the thalamus enhances cue-induced reinstatement in goal-trackers, but not sign-trackers. Psychopharmacology (Berl) 2018;235:999–1014. 10.1007/s00213-017-4816-1.

[56] Robbins TW, Everitt BJ. Neurobehavioural mechanisms of reward and motivation. Curr Opin Neurobiol 1996;6:228–36. 10.1016/S0959-4388(96)80077-8.

[57] White FJ, Kalivas PW. Neuroadaptations involved in amphetamine and cocaine addiction. Drug Alcohol Depend 1998;51:141–53. 10.1016/S0376-8716(98)00072-6.

[58] Hearing M. Prefrontal-accumbens opioid plasticity: Implications for relapse and dependence. Pharmacol Res 2019;139:158–65. 10.1016/j.phrs.2018.11.012.

[59] McGinty JF, Otis JM. Heterogeneity in the Paraventricular Thalamus: The Traffic Light of Motivated Behaviors. Front Behav Neurosci 2020;14.

[60] Gao C, Leng Y, Ma J, Rooke V, Rodriguez-Gonzalez S, Ramakrishnan C, Deisseroth K, Penzo MA. Two genetically, anatomically and functionally distinct cell types segregate across anteroposterior axis of paraventricular thalamus. Nat Neurosci 2020;23:217–28. 10.1038/s41593-019-0572-3.

[61] Penzo MA, Gao C. The paraventricular nucleus of the thalamus: an integrative node underlying homeostatic behavior. Trends Neurosci 2021;44:538–49. 10.1016/j.tins.2021.03.001.

[62] Mount KA, Kuhn HM, Hwang E-K, Beutler MM, Wolf ME. Incubation of oxycodone craving is associated with CP-AMPAR upregulation in D1 and A2a receptor-expressing medium spiny neurons in nucleus accumbens core and shell. Neuropharmacology 2026;287:110816. 10.1016/j.neuropharm.2025.110816.

